# A limbic circuit selectively linking active escape to food suppression

**DOI:** 10.1101/683946

**Authors:** Estefania P. Azevedo, Bowen Tan, Lisa E. Pomeranz, Violet Ivan, Robert N. Fetcho, Marc Schneeberger, Katherine R. Doerig, Conor Liston, Jeffrey M. Friedman, Sarah A. Stern

## Abstract

Stress and anxiety are precipitating factors for eating disorders, but the neural basis linking stress to alterations in feeding is not well understood. Here we describe a novel population of stress-responsive neurons in the lateral septum (LS) of mice that express neurotensin (LS^NTS^) in a sexually dimorphic, estrous cycle-dependent manner. We used in vivo imaging to show that LS^NTS^ neurons are activated by stressful experiences when flight is a viable option, but not by a stressful experience associated with freezing or immobility. LS^NTS^ activation leads to a decrease of food intake and body weight in mice, without altering locomotion or other behaviors associated with anxiety. Molecular profiling of LS^NTS^ neurons showed that these neurons co-express Glp1r (glucagon-like peptide 1 receptor), and both pharmacologic and genetic manipulations of Glp1r signaling in the LS recapitulates the behavioral effects of LS^NTS^ activation. Finally, we mapped the outputs of LS^NTS^ neurons and show that activation of LS^NTS^ nerve terminals in the lateral hypothalamus (LH), a well-established feeding center, also decrease food intake. Taken together, these results show that LS^NTS^ neurons link stress and anorexia via effects on hypothalamic pathways regulating food intake.

## INTRODUCTION

A key function of the central nervous system (CNS) is to sense external conditions and activate an appropriate behavioral response (McEwen and Akil, 2020). The detection of stressful conditions, in particular, is of critical importance by enabling animals to remove themselves from dangerous situations. These responses are also associated with other adaptive behavioral effects including reduced feeding until the period of danger has passed, and these stereotyped responses are conserved across organisms ranging from flies to mice and humans (Yau and Potenza, 2013; Surendran et al., 2017).

We therefore set out to map neural circuits linking stress to feeding by initially studying acute restraint stress, a behavioral paradigm in rodents which leads to a robust decrease in food intake for up to 48 hours (Donohoe, 1984; Shimizu et al., 1989). Previous studies have shown increased expression of c-fos, an early immediate gene, in many brain areas following stress, including regions of the limbic system (e.g. lateral septum, Sheehan et al., 2004). However, the cell-type specificity and the activation dynamics of stress-responsive neurons in specific regions of the limbic system has not been determined, nor has the mechanism by which these neurons regulate feeding.

In this study, we used an unbiased transcriptomic approach (Knight et al., 2012) to map and identify neurons in the limbic system that are activated during stress-induced anorexia. Initial studies revealed that neurons in the lateral septum that express neurotensin are acutely activated by restraint stress and we found that acute or chronic chemogenetic activation of these neurons leads to significant reductions of food intake and body weight. Chronic stress is also a risk factor for the development of many psychiatric diseases that are characterized by maladaptive responses (Yau and Potenza, 2013), including eating disorders, which have a significantly greater incidence in females (Guarda et al., 2015). Accordingly, we also found that the activity of these neurons is regulated by the levels of female sex hormones, raising the possibility that LS^NTS^ neurons could provide an entry point for studying neural circuits that may contribute to eating disorders.

## RESULTS

### Identification of a limbic population activated by acute restraint stress

In order to identify specific neuronal populations that link stress to reduced food intake, we first measured food intake in wild-type C57Bl/6J mice after one hour of acute restraint stress and mapped the expression pattern of c-fos throughout the whole mouse brain. As previously reported, acute restraint stress for a period of 1h led to a highly significant decrease in food intake for the ensuing 2 hours (Fig. 1A, Two-way ANOVA with Bonferroni post-hoc, *p<0.05, ***p<0.001). One hour of restraint stress also led to increased numbers of c-fos+ cells in a number of brain regions within the limbic system: the basolateral amygdala [BLA], bed nucleus of the stria terminalis [BNST], cortex (insular cortex [IC]) (Figure 1B and Table S1) and elsewhere. Consistent with previous studies (Kubo et al., 2002), acute restraint stress led to a marked increase in the number of c-fos+ cells in the LS compared to controls (Figure S1A, Unpaired Student’s t-test, *p<0.05). The LS is part of the anterior limbic system and is known to regulate several emotional behaviors, particularly stress, anxiety and aggression (Kubo et al., 2002; Anthony et al., 2014; Wong et al., 2016). In addition, previous studies from our group and another showed that the artificial activation of neurons in the LS has a negative effect on feeding (Azevedo et al., 2019; Sweeney and Yang, 2016), raising the possibility that specific neurons in this region might link restraint stress to a non-homeostatic reduction in food intake.

**Figure 1.**
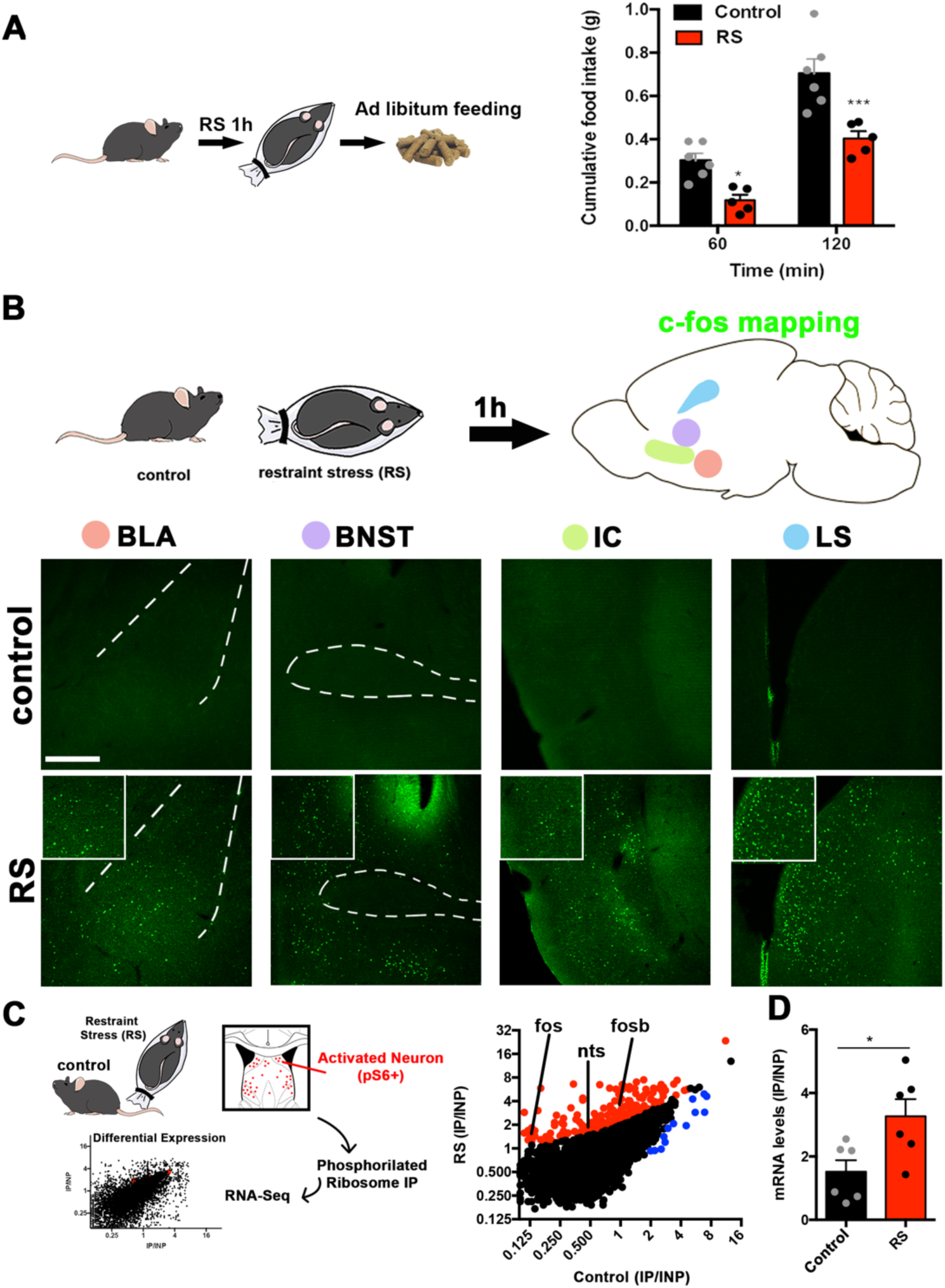
Identification of lateral septum neurons activated by acute restraint stress. (A) Left panel, experimental scheme. Right panel, food intake, measured in grams (g), in control (black) and RS (red) mice after 60 and 120min following RS. n=5(RS), n=6(control), Two-way ANOVA with post-hoc Bonferroni correction, Time: F(1,9)=175.9, p<0.0001; Subject: F(1,9)=16.86, p=0.003; Interaction: F(1,9)=4.975, p=0.0527). (B) Activity mapping in the whole brain using c-fos staining in control (naïve) and restraint stressed (RS) mice. Areas with the highest c-fos expression (green) difference are shown: basolateral amygdala (BLA), bed nucleus of the stria terminals (BNST), insular cortex (IC) and lateral septum (LS). Insets show 2x digital zoom of c-fos expression. Scale bars, 50 μm. (C) Activity-based transcriptomics (PhosphoTrap) of the lateral septum of control (naïve) and restraint stressed mice. Middle panel, plot depicting the average IP/INP value (log2) of all genes analyzed in control and restraint stress (RS) samples. Enriched genes (>2-fold; red) and depleted genes (<2-fold; blue) are shown, and activation or target gene markers are shown: Nts (neurotensin), fos and fosb (n = 2). All genes depicted have q-values <0.05 as calculated by Cufflinks. (D) mRNA levels of Nts evaluated using Taqman qPCR in control and restraint stress (RS) samples: control (black bars) and RS (red bars) are compared (n = 6). Unpaired Student’s t-test, *p < 0.05, t=2.692, df=10. Data are represented as mean ± SEM.

We next set out to determine the molecular identity of the neurons in the LS that are activated by acute restraint stress using PhosphoTrap, an unbiased transcriptomic method to molecularly profile activated neurons (Knight el al., 2012). This method takes advantage of the fact that neuronal activation results in a cascade of signaling events culminating in the phosphorylation of the ribosomal protein S6 protein (pS6). These phosphorylated ribosomes can be immunoprecipitated from mouse brain homogenates, thereby enriching mRNAs selectively expressed in the activated neuronal population (Fig. 1C, left panel). After polysome immunoprecipitation and RNA extraction and sequencing, RNAs enriched relative to total RNA have been shown to mark activated neurons (Knight et al., 2012; Azevedo et al., 2014; Stern et al., 2019). We applied this method to identify markers for neurons in the lateral septum that are activated after acute restraint stress relative to control mice (naïve) (Fig. 1C). The enrichment for each gene was calculated as the number of reads in the immunoprecipitated RNA (IP) relative to the total input RNA (IP/INP; Fig. 1C). As expected based on data from immediate early gene mapping, we found an enrichment for activity-related genes (*FosB* and *Fos*) in the samples from mice exposed to acute restraint stress, (see Fig. 1B, LS and Figure S1A). We next analyzed the data to identify other genes that were significantly enriched (>2-fold or greater and q-value <0.05) in the samples from mice exposed to acute restraint stress (RS) compared to naïve mice (q value <0.05, Fig. 1C and Table S2). We found enrichment for several markers, including the gene for the neuropeptide neurotensin (*Nts*, 2.41-fold increase compared to naïve samples). We confirmed the enrichment using qPCR and found a 3.6-fold enrichment in neurotensin mRNA in the samples from the lateral septum of mice exposed to acute restraint stress compared to samples from naive mice (Unpaired Student’s t-test, p<0.05).

### LS^NTS^ neurons respond to stress in a sexual dimorphic manner

We further confirmed the activation of Nts-expressing neurons by acute restraint stress by analyzing the colocalization of *Nts* and *Fos* mRNA expression in the LS using *in situ* hybridization (Fig. 2). Both male and female mice showed significant colocalization between *Fos* and *Nts* in the LS after restraint stress compared to their control non-stressed, naïve counterparts (∼65 cells in stressed females vs. 2 cells in non-stressed females; ∼37 cells in stressed males vs. 22 cells in non-stressed males; **p<0.01, *p<0.05, Fig. 2B and C). Because the effects of acute stress are modulated by sex hormones in particular in eating disorders, we analyzed the co-expression of *Nts* and *Fos* in stressed female mice during different phases of the estrous cycle. Female mice in estrus exposed to acute restraint stress had a ∼2-fold higher levels of *Nts* and *Fos* colocalization compared to male mice (Figure 2B and C). Furthermore, the colocalization of *Nts* and *Fos* was significantly higher in female mice during the estrus phase compared to those that were in diestrus phase (Figure 2B, ∼ 60 cells/slice in estrus compared to ∼30 cells/slice in diestrus). The extent of the colocalization of *Nts* and *Fos* after acute restraint stress (RS) was similar in females during the diestrus phase to males (Figure 2B and C, ∼37 cells/slice in males and ∼25 cells/slice in females in diestrus). We also found that the higher levels of *Fos* colocalization with *Nts* in stressed female and male mice was due to both an increase in the levels of *Fos* in *Nts* neurons and a concomitant induction of *Nts* mRNA during stress (Figure S1B and C). In contrast, neurotensin mRNA was almost completely absent in the LS in non-stressed female controls (Fig 1B). These results suggest that that activity of Nts neurons as well as the expression of the Nts gene in the LS is regulated by acute restraint stress in a sexually dimorphic manner and potentiated by the increased levels of female sex hormones during estrus.

**Figure 2.**
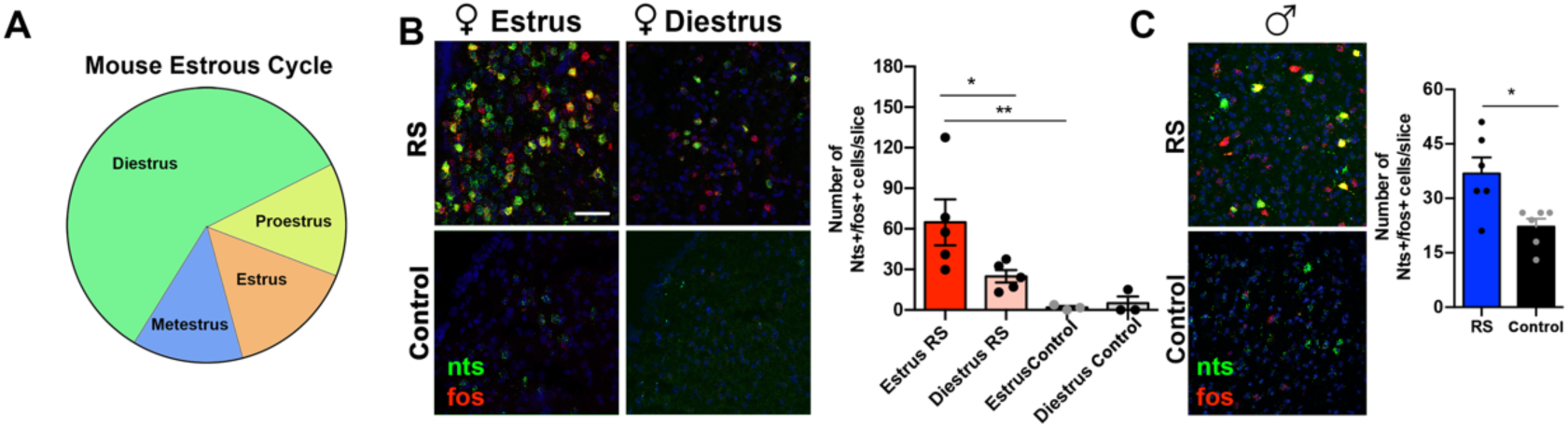
Nts expression is regulated by stress and the estrous cycle in females. (A) The mouse estrous cycle: the different phases of the mouse estrous cycle (diestrus, proestrus, estrus, metestrus) are shown in different colors. (B) In situ hybridization showing the expression of Nts (green) and fos (red) mRNA in females in estrus and diestrus exposed to acute restraint stress (RS) or naïve (control). Quantification of the number of Nts+/fos+ cells per slice/mice are shown, n=3-5, Nested one-way ANOVA, F=6.366, DFn=3, Dfd=12. (C) In situ hybridization showing the expression of Nts (green) and fos (red) mRNA in males exposed to acute restraint stress (RS) or naïve (control). Quantification of the number of Nts+/fos+ cells per slice/mice are shown, n=6, Unpaired Student’s t-test, *p<0.05, t=2.962, df=10. Data are represented as mean ± SEM.

### LS^NTS^ neurons regulate food intake and body weight

We next tested whether chemogenetic activation of Nts neurons would lead to a concomitant decrease in food consumption. We injected an AAV encoding a Cre dependent excitatory DREADD, (DIO) hM3Dq, a Gq-coupled (designer receptor exclusively activated by designer drugs, Armbruster et al., 2007), into the LS of Nts-Cre mice. Injection of clozapine-N-oxide (CNO) into these mice increased c-fos in the LS (Figure S1D, Unpaired Student’s t-test) and significantly decreased food intake compared to mice treated with CNO after injection of a control Cre-dependent mCherry AAV into Nts-cre mice (Fig. 3A, red bars vs black bars, Two-way ANOVA with Bonferroni post-hoc, *p<0.05, ***p<0.001) as well as saline-injected mice expressing the DREADD (Figure S1E). This robust decrease in food intake was evident after 1 hour and persisted for 4, 8 and 24 hours after a single CNO injection (Figure S1F). Chronic stimulation of Nts neurons for 3 days in the LS also significantly decreased food intake (Fig. 3B, Two-way ANOVA with Bonferroni post-hoc, **p<0.01, ****p<0.0001) and significantly reduced body weight relative to the aforementioned controls (Fig. 3C, Unpaired Student’s t-test, p<0.05).

**Figure 3.**
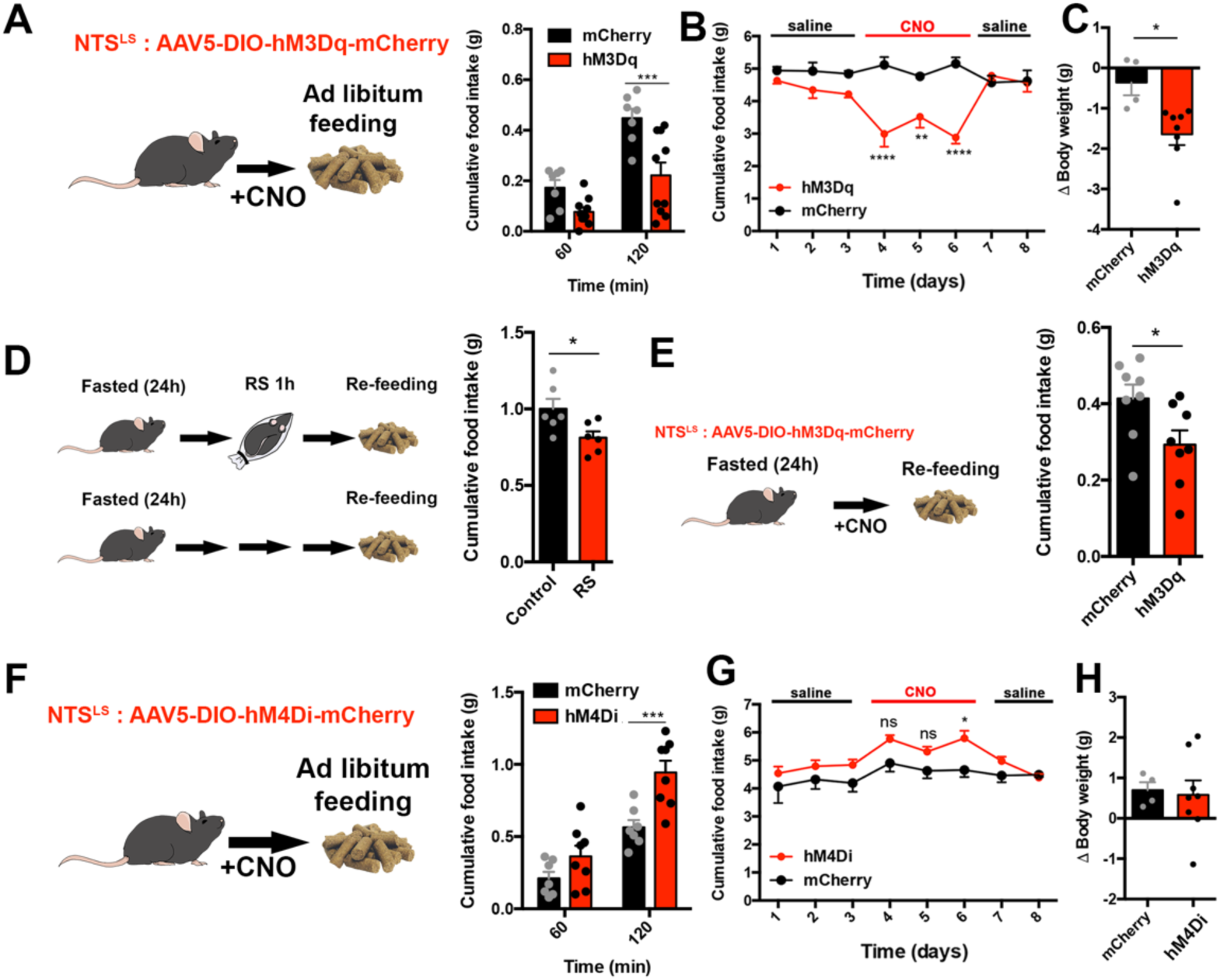
LS^NTS^ neurons regulate food intake and body weight in mice. (A) Left panel, experimental scheme. NTS-cre mice were injected with an AAV encoding the hM3Dq activatory DREADD receptor in the LS. Middle panel, cumulative food intake, measured in grams (g) of control mCherry (black) and hM3Dq (red) expressing NTS-cre mice after 60 and 120 min following CNO injection. n=7(mCherry), n=10(hM3Dq), Two-way ANOVA with post-hoc Bonferroni correction, Time: F(1,15)=55.46, p<0.0001; Subject: F(1,15)=12.84, p=0.0027; Interaction: F(1,15)=5.45, p=0.034). (B) Daily food intake, measured in grams (g) daily during injections of saline (days 1-3, 7-8) or CNO (days 4-6) in control mCherry (black) or hM3Dq (red) expressing NTS-cre mice. n=8; Two-way ANOVA with post-hoc Bonferroni correction, Time: F(7,72)=2.871, p=0.01; Subject: F(1,72)=46.92, p<0.0001; Interaction: F(7,72)=6.48, p<0.0001). (C) Body weight (delta), measured in grams (g) after 3 days of daily CNO injections in control mCherry (black) or hM3Dq (red) expressing NTS-cre mice. n=8; Unpaired Student’s t-test, *p<0.05, t=2.903, df=10. (D) Left panel, experimental scheme. Right panel, cumulative food intake, measured in grams (g), in control naïve mice (control, black) or mice that underwent restraint stress (RS, red), measured 2h after re-feeding after an overnight fast, n=6. Unpaired Students’ t-test: T(10)=2.427, p=0.0356. (E) Left panel, experimental scheme. Right panel, cumulative food intake, measured in grams (g), in control mCherry (black) or hM3Dq(red) expressing NTS-cre mice, measured 2h following re-feeding after an overnight fast, n=8, Paired Student’s t-test, *p<0.05, t= 3.171, df=7. (F) Left panel, experimental scheme. Right panel, cumulative food intake, measured in grams (g), in control mCherry (black) and hM4Di (red) expressing NTS-cre mice after 60 min and 120 min following saline injection. n=7(mCherry), n=8(hM4Di). Two-way ANOVA with post-hoc Bonferroni correction, Time: F(1,19)=415.8, p<0.0001; Subject: F(1,13)=1.042, p=0.326; Interaction: F(4,52)=2.839, p=0.0333. (G) Daily food intake, measured in grams (g) after daily injections of saline (days 1-3, 7-8) or CNO (days 4-6) in control mCherry (black) or hM4Di (red) expressing NTS-cre mice. n=7; Two-way ANOVA with post-hoc Bonferroni correction, Time: F(7,96)=3.949, p=0.0008; Subject: F(1,96)=18.95, p<0.0001; Interaction: F(7,96)=0.876, p=0.5285). (H) Body weight (delta), measured in grams (g) after 3 days of daily CNO injections in control mCherry (black) or hM4Di (red) expressing NTS-cre mice. n=8; Unpaired Student’s t-test, p=0.83, t=0.2168, df=10. Data are represented as mean +- SEM.

We next tested whether restraint stress could override homeostatic feeding in animals that were provided with food after an overnight fast. We found that fasted mice that had also been exposed to restraint stress for 1 hour consumed significantly less food during refeeding than controls (Fig. 3D, Unpaired Student’s t-test, *p<0.05), indicating that acute stress can override the hyperphagia that follows a fast. Consistent with this, chemogenetic activation of Nts neurons in the LS in mice also suppressed food intake in mice that were refed after that a 24h fast (Fig. 3E, Unpaired Student’s t-test, *p<0.05).

To investigate whether inactivation of Nts neurons could have an opposite effect in food intake and stimulate consumption, we injected Cre conditional AAV expressing the inhibitory hM4Di receptor or a control AAV expressing mCherry into the LS of Nts-cre mice. Acute inhibition of LS^NTS^ neurons led to an increase in food intake after CNO injection compared to mCherry controls in the first 2hrs (Fig. 3F, Two-way ANOVA with Bonferroni post-hoc, ***p<0.001) and at 8 and 24h (Figure S1H). As expected, food intake was unchanged after saline-injection into mice expressing the inhibitory DREADD (Figure S1G). Chronic inhibition of Nts neurons in the LS for 3 days produced an increase in food intake that was significant in the 3^rd^ day (Fig. 3G, Two-way ANOVA with Bonferroni post-hoc, *p<0.05) but did not increase body weight (Fig. 3H, Unpaired Student’s t-test, p=0.83). These results suggest that inhibition of Nts neurons increases food intake.

Interestingly, LS^NTS^ chemogenetic activation did not alter behavior during open field thigmotaxis, general locomotion (Figure S2A-C), when animals were in an open-field with novelty (Figure S2D), elevated plus maze (Figure S2E-G), or during novelty-suppressed feeding (Figure S2H) Thus, the data show LS^NTS^ neurons do not appear to play a discernible role in the other anxiogenic behaviors that were tested and appear to be specific for feeding.

### LS^NTS^ neurons are selectively tuned to active escape

We next set out to assay the dynamics of LS^NTS^ activation in response to stress using in vivo calcium imaging. We injected AAVs expressing a Cre-dependent GCaMP6 into the LS of NTS-Cre mice and recorded activity using *in vivo* fiber photometry in freely behaving animals while time-locking behavior with changes in calcium signals (Gunaydin et al., 2014;Fig. 4 and Figure S3A). GCaMP6s expression and optic fiber placement were confirmed by immunohistochemistry following the experiments (Figure S3B). We first assessed activity during acute restraint stress by manually restraining mice for a 1-minute period. Based on the c-fos data (Fig. 1), we expected to see an increase in calcium activity while the animal was being actively restrained. Surprisingly, we did not see a change in the calcium signal when the animal was immobile. Rather, we only found increases in the calcium signal when animals were actively struggling to escape immediately before and after the manual restraint. This reduced neural activity was seen either when the animals were completely immobilized or when they had ceased struggling (Fig. 4A, one-way ANOVA, multiple comparisons with Bonferroni post-hoc, *p<0.05). We then recorded LS^NTS^ calcium signals in animals that were exposed to a one-minute tail suspension test during which time the animals actively try to escape and, consistent with the previous results, we also observed activation of LS^NTS^ neurons while the animals struggled (Fig. 4B, Paired Student’s t-test, *p<0.05).

**Figure 4.**
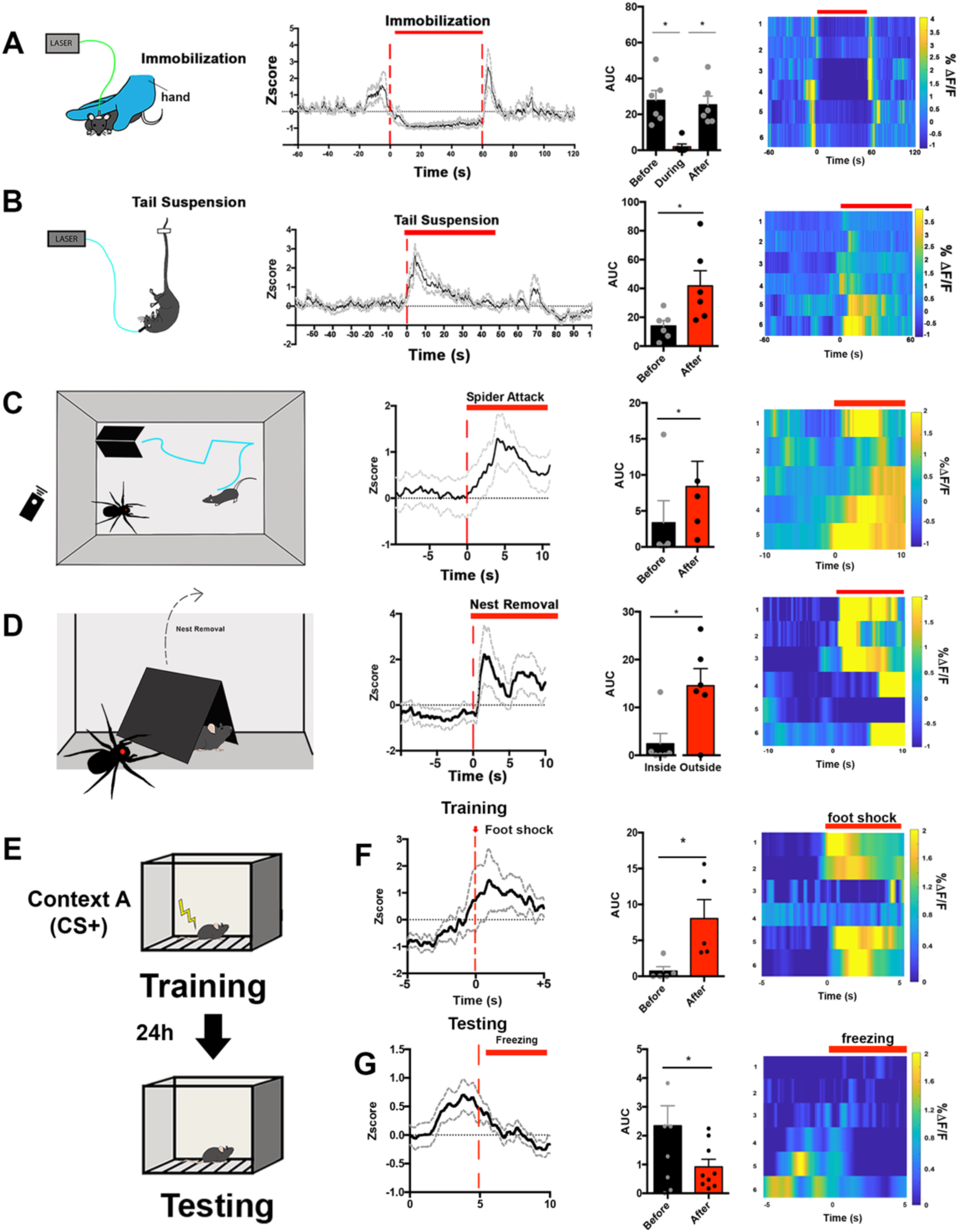
LS^NTS^ neurons are specifically activated by stressful situation involving active coping. (A) Left panel, experimental scheme: mice were manually immobilized for the period of 1 min. Ca^+2^ activity was recorded before, during and after the period of manual immobilization. Middle, Average GCaMP6s z-score (black lines) and SEM (gray dotted lines) across all recordings and time-locked to immobilization start. Right middle panel, quantification of the area under the curve of Ca^+2^ transients before, during and after immobilization. One-way ANOVA with repeated measures and post-hoc Bonferroni correction, F(1.833, 9.165)=9.888, p=0.0057. Right panel, heat maps represent the average % ΔF/F from each gCaMP6s recordings time-locked to immobilization start. n=6. (B) Left panel, experimental scheme: mice were tail suspended for the period of 1 min. Ca^+2^ activity was recorded before, during and after the period of tail suspension. Middle, Average GCaMP6s z-score (black lines) and SEM (gray dotted lines) across all recordings and time-locked to tail suspension start. Right middle panel, quantification of the area under the curve of Ca^+2^ transients before, during and after tail suspension. Paired Student’s t-test, t=2.659, df=5. Right panel, heat maps represent the average % ΔF/F from each gCaMP6s recordings time-locked to tail suspension start. n=6. (C) Left panel, experimental scheme: mice were exposed to an open field arena containing a robotic, remote-controlled spider predator and a designated safe “nest” (black triangular hut). Ca^+2^ activity was recorded during the whole session. Middle, Average GCaMP6s z-score (black lines) and SEM (gray dotted lines) across all recordings and time-locked to the robotic spider attack. Right middle panel, quantification of the area under the curve of Ca^+2^ transients before and after the robotic spider attack. Paired Student’s t-test, t=3.507, df=4. Right panel, heat maps represent the average % ΔF/F from each gCaMP6s recordings time-locked to the robotic spider attack. n=6. (D) Left panel, experimental scheme: the designated safe “nest” (black triangular hut) was removed and mice that had been attacked by a robotic spider predator was exposed to the open field containing the spider. Ca^+2^ activity was recorded during the whole session. Middle, Average GCaMP6s z-score (black lines) and SEM (gray dotted lines) across all recordings and time-locked to the removal of the nest. Right middle panel, quantification of the area under the curve of Ca^+2^ transients before and after the removal of the nest. Paired Student’s t-test, t=3.305, df=5. Right panel, heat maps represent the average % ΔF/F from each gCaMP6s recordings time-locked to the removal of the nest. n=6. (E) Experimental scheme for contextual fear conditioning: mice were exposed to an operant chamber where they received a mild foot shock (training) and after 24h mice were re-exposed to the operant chamber (testing). Ca^+2^ activity was recorded during training and testing. (F) First panel, average GCaMP6s z-score (black lines) and SEM (gray dotted lines) across all recordings and time-locked to the foot shock received on the training day. Second panel, quantification of the area under the curve of Ca^+2^ transients before and after the foot shock. Paired Student’s t-test, t=3.230, df=4. Third panel, heat maps represent the average % ΔF/F from each gCaMP6s recordings time-locked to the freezing behavior. n=6. (G) First panel, average GCaMP6s z-score (black lines) and SEM (gray dotted lines) across all recordings and time-locked to freezing behavior on the testing day. Second panel, quantification of the area under the curve of Ca^+2^ transients before and after the freezing behavior. Paired Student’s t-test, t=2.833, df=8. Third panel, heat maps represent the average % ΔF/F from each gCaMP6s recordings time-locked to the freezing behavior. n=6. Data are represented as mean +-SEM.

We thus considered the possibility that LS^NTS^ neurons are specifically activated by stressful situations where the animal has an ability to escape i.e; stress-induced flight. To test this further, we exposed mice to a simulated “predator” using a remote-controlled robotic spider (Fig. 4C, left panel). In this paradigm, mice are habituated to an open field arena containing a homemade opaque acrylic nest that serves as a protected “safe zone”. The remote-controlled robotic spider was placed into the arena and after 5 min of habituation the spider movement was manually controlled so that it only moved when mice left the acrylic nest. We observed that forward movement of the robotic spider toward to the mouse induced a rapid retreat leading the mice to immediately return to the inside the acrylic nest (Supplementary Video 1). We simultaneously used *in vivo* fiber photometry to monitor the activity of LS^NTS^ neurons during this simulated predator attack and found that movement of the robotic spider towards the mouse led to a strong activation of LS^NTS^ neurons that persisted until the mouse returned to the nest (Fig. 4C, Paired Student’s t-test, *p<0.05). Similarly, we observed a rapid and sustained increase in the calcium signal when the nest was removed and mice were constantly exposed to the robotic spider (Fig. 4D, Paired Student’s t-test,* p<0.05). In aggregate, these results show that LS^NTS^ neurons are activated by stressful stimuli in which the animal’s response includes active movement such as struggling or flight (active coping strategies) and further suggest that they are not activated by stressful stimuli associated with immobility or freezing i.e; passive coping responses.

To further confirm the tuning of LS^NTS^ to stress stimuli that involved active coping, we monitored the activity of LS^NTS^ neurons in mice during a contextual fear conditioning protocol (CFC), in which mice are trained to associate a novel environment with an unpredicted foot shock. This task enables a comparison of different types of stress responses because during the training phase, the foot shock leads to active movement away from the shock source, while during testing, exposure to the contextual cue results in freezing behavior (Figure S3G and H). Moreover, unlike restraint, which is imposed by the experimenter, freezing is self-initiated. Consistent with the prior results, we found that during the training session, LS^NTS^ calcium signals were increased following the foot shock, time-locked with movement, as the mice jumped away from the shock (Fig. 4E, Paired Student’s t-test, p<0.05). However, during the recall test the following day, when mice exhibited freezing behavior after being placed in the chamber that they associated with a foot shock, the LS^NTS^ calcium signal was significantly decreased compared to the period before the onset of freezing or compared to the signal during the active movement exhibited during the foot shock (Fig. 4F, Paired Student’s t-test, p<0.05). Taken together, these data are consistent with the possibility that the activity of these neurons is tuned to active, but not passive, responses to stressful stimuli. They also indicate that these neurons remain active until the animal is in an environment that is either perceived to be “safe” or until the animal freezes as part of a passive coping strategy. In contrast, we did not find any significant temporal correlation between calcium signals in LS^NTS^ neurons and feeding (Figure S3C and D) or voluntary movement in an open field arena (Figure S3E and F).

### Molecular Profiling of LS^NTS^ neurons reveal the expression of a diversity of molecules and receptors

The majority of the neurons in the LS are GABAergic and a number of neuropeptides and specific receptors are expressed in this region in addition to neurotensin (Lein et al., 2007). In order to identify possible inputs that regulate LS^Nts^ neurons, we molecularly profiled LS^NTS^ neurons by injecting a Cre-dependent adeno-associated virus expressing an GFP-tagged ribosomal protein (AAV-Introvert-EGFPL10a) into the LS of NTS-Cre mice (Nectow et al., 2017). Polysomes were immunoprecipitated with an anti-GFP antibody to profile LS^NTS^ neurons using viral-based translating ribosome affinity purification followed by RNA-seq (viralTRAP; Fig. 5A). The enrichment for each gene was calculated as the number of reads in the immunoprecipitated RNA (IP) relative to the total input RNA (IP/INP; Fig 5B). Several genes were significantly enriched in the immunoprecipitated fraction (IP) including Gfp, somatostatin Sst, as well as Glp1r, Cartpt and Mc3r (Fig. 5B and C, Table S3). We confirmed the co-expression of these transcripts with Nts using qPCR of the precipitated RNAs (Fig. 5C) and *in situ* hybridization which showed that ∼80% of septal Nts neurons co-express Slc32a1 (vgat), a marker of GABAergic neurons, (Figure 5D, Unpaired Students’ t-test, **p<0.01). We also verified co-expression of Nts with Glp1r, Sst, Cartpt and Mc3r with 70% of LS^NTS^ neurons expressing Glp1r, 50% expressing Sst, 40% expressing Cartpt and 20% expressing Mc3r (Fig. 5E-I).

**Figure 5.**
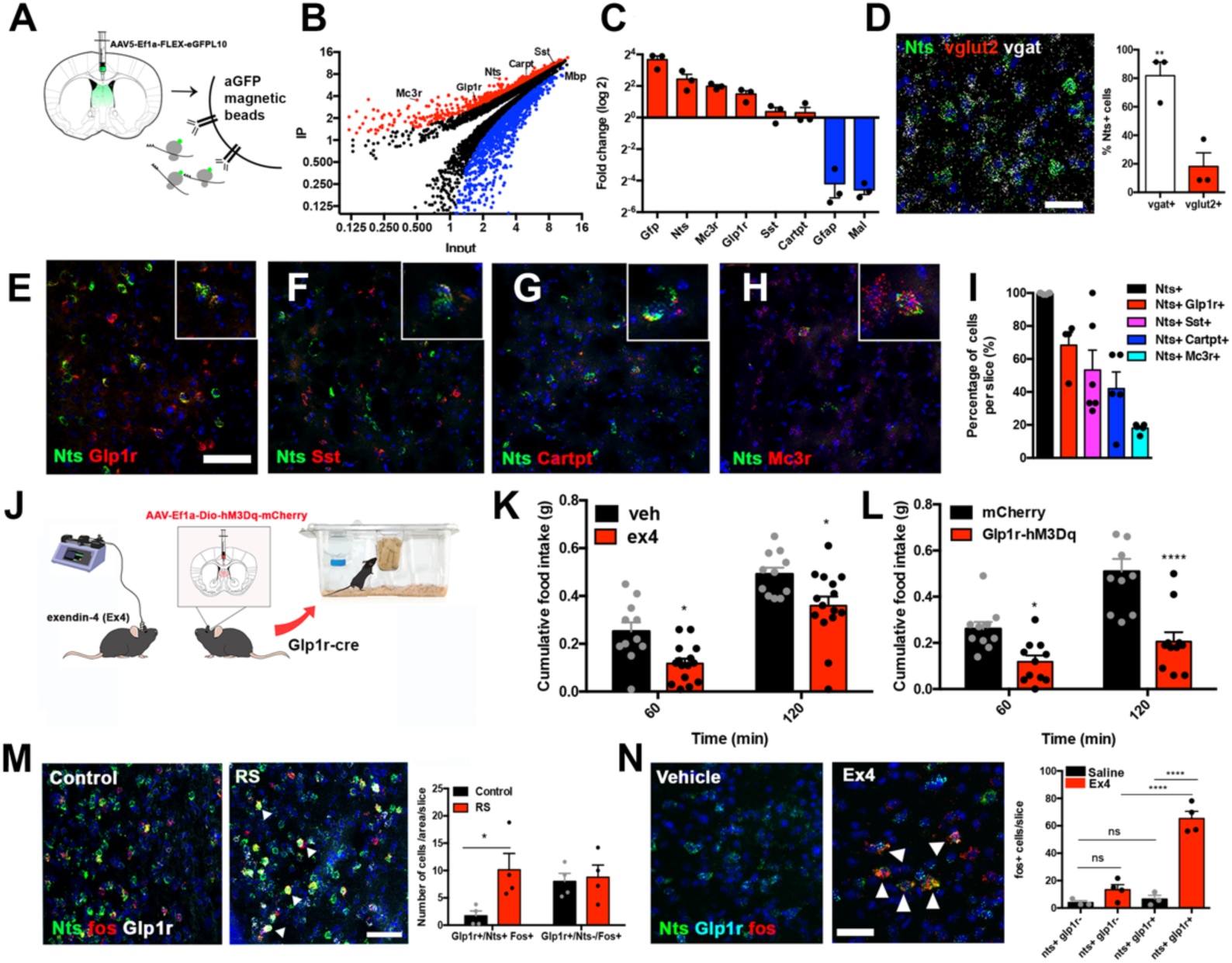
Molecular and functional profiling of LS^NTS^ neurons. (A) Experimental scheme of the viralTrap experiment. NTS-cre mice are injected with an AAV expressing cre-dependent GFPL10 (AAV-Ef1a-FLEX-eGFPL10) into the LS and then GFP+ polysomes are immunoprecipitated. (B) Plot depicting the average IP value and average input values (log2) of all genes analyzed. Enriched genes (>2-fold; red) and depleted genes (<2-fold; blue) are shown (q<0.05), and selected markers are shown: mc3r, Glp1r, Nts, Sst, Cartpt and Mbp (n = 3). All genes depicted have q-values <0.05 as calculated by Cufflinks. (C) Average fold of change (IP/INP Log2) assessed by Taqman qPCR of enriched positive control markers Gfp and Nts, significantly enriched genes Mc3r, Glp1r, Sst and Cartpt (red) and depleted negative control markers Gfap and Mal (blue) (n = 3 biological replicates). (D) Left panel, representative in situ hybridization image of Nts (green), vglut2 (red), vgat (white) and DAPI (blue). Right panel, quantification of Nts cells which express vgat (white) and vglut2 (red). n=3, Unpaired Students’ t-test p=0.009; t(4)=4.791. Scale bars, 50 um. (E) Representative in situ hybridization image of Nts (green) and Glp1r (red). 2x digital zoom is presented in the inset. Scale bars for panels e-h, 50 um. (F) Representative in situ hybridization image of Nts (green) and Sst (red). Closer magnification is inset. 2x digital zoom is presented in the inset. (G) Representative in situ hybridization image of Nts (green) and Cartpt (red). 2x digital zoom is presented in the inset. (H) Representative in situ hybridization image of Nts (green) and Mc3r (red). 2x digital zoom is presented in the inset. (I) Quantification of the percentage of Nts cells (black) expressing Nts/Glp1r (red), Sst (magenta), Cartpt (blue) and Mc3r (cyan). n=3(Nts), n=4(Nts/Glp1r, Nts/Mc3r), n=6(Nts/Sst), n=5(Nts/Cartpt) (J) Left panel, experimental scheme. Wild-type mice are injected with exendin-4 (Ex4) directly into the LS or Glp1r-cre mice with viral expression of hM3Dq activatory DREADD receptor are tested for cumulative food intake. (K) Cumulative food intake measured in grams (g) of mice following Vehicle (black) and Ex4 (red) injection in the LS. n=11(Vehicle), n=15(Ex4), Two-way ANOVA with post-hoc Bonferroni correction (Time: F(1,24)=100.8, p<0.0001; Treatment: F(1,24)=11.54, p=0.0024; Interaction: F(1,24)=0.002; p=0.963. (L) Cumulative food intake measured in grams (g) of control mCherry (black) and hM3Dq (red) expressing Glp1r-cre mice following CNO injection. Two-way ANOVA with post-hoc Bonferroni correction. n=10(mCherry), n=11(hM3Dq). (Time: F(1,19)=40,67, p<0.0001; Subject: F(1,19)=20.68, p=0.0002; Interaction: F(1,19)=9,407, p=0.0063. (M) Representative images of Nts (green), fos (red) and Glp1r (white) in control and restraint stress (RS) mice. Right, quantification of co-localization between Nts, fos and Glp1r+ cells in control (black) and RS (red) mice. n= 4, Two-way ANOVA with post-hoc Bonferroni correction, *p<0.05, Row factor: F(1,12)=1.460, p=0.2502, Column Factor: F(1,12)=5.056, p=0.0441, Interaction: F(1.12)=3.494, p=0.0862. Scale bars, 50 um. (N) Left, representative in situ hybridization image of Nts (green), Glp1r (blue) and Fos (red) of mice injected with vehicle or the Glp1r agonist Exendin-4 (Ex4). White arrows point to cells co-expressing Nts, Glp1r and Fos. Right, quantification of Fos+ cells expressing Nts and/or Glp1r in saline (black) or Ex4 (red) injected mice. n=3(Control), n=4 (Ex4); Two-way ANOVA with Bonferroni correction (Subject: F(1,10)=71.7, p<0.0001; Treatment: F(1,10)=45,88, p=0.0001; Interaction: F(1,10)=37.58, p<0.0001). Data are represented as mean +- SEM.

Many of the genes co-expressed with Nts in LS^NTS^ neurons are known to regulate feeding behaviors in other brain areas, including Cartpt (Kristensen et al., 1998), Glp1r (Kanoski et al., 2016), Mc3r (Marks et al., 2006) and Sst (Luo et al., 2018). For this reason, we asked whether activation of neurons expressing these genes could replicate the effects of LS^NTS^ neuronal activation. Similar to chemogenetic activation of LS^NTS^ neurons, activation of Glp1r neurons within the LS with the Glp1r agonist exendin-4 (Fig.5J) resulted in a robust decrease in food intake after 2, 12 and 24h (Figure S4A-C). In addition, chemogenetic activation of Glp1r neurons by injecting the activatory hM3Dq DREADD into the LS of Glp1R-cre mice also reduced food intake (Two-way ANOVA with Bonferroni post-hoc, *p<0.05, ****p<0.0001). Neither control mice expressing the DREADD but injected with saline nor mice injected with mCherry-expressing AAVs and given CNO exhibited any change in food intake (Fig. 5K and L; Figure S4A-C). In contrast, food intake was unchanged after infusion of the Mc3r agonist (gamma-melanocyte-stimulating hormone; y-msh) into the septum or activation of Cartpt or Sst neurons by injecting an activatory DREADD (hM3Dq) into Cartpt-cre and Sst-cre animals (Figure S4D-F). Activation of these other LS^NTS^ subpopulations (Figure S4G-I) also had no effect on anxiety as assessed using an elevated plus maze (Figure S4J-L).

Glp-1 signaling has been shown to elicit an anorectic effect and blocking of Glp1r signaling using specific antagonists has been shown to blunt stress-induced anorexia (Terril et al., 2018). After acute restraint stress, we observed significantly increased expression of *Fos* only in LS^NTS^ neurons that also co-express *Glp1r*, while Fos expression was unchanged in *Glp1r* neurons that do not express *Nts* (Fig. 5M, Two-way ANOVA with Bonferroni post-hoc, *p<0.01). Furthermore, infusion of Exendin-4 (ex4), a Glp1r agonist, into the LS increased *Fos* expression in ∼65% of LS^NTS^ neurons co-expressing *Glp1r* (Fig. 5N, Two-way ANOVA with Bonferroni post-hoc, ***p<0.0001), but not in *Nts* neurons that do not co-express *Glp1r* (Fig. 5K). These results identify a stress-responsive subpopulation of LS^NTS^ neurons co-expressing Glp1r that negatively regulates food intake and further shows that Glp-1 (glucagon-like peptide-1) signaling to this subpopulation of LS^NTS^ neurons regulates their activity.

### Mapping the downstream projections of LS^NTS^ neurons

Finally, we set out to map the functional axonal projections of LS^NTS^ neurons by injecting a cre-dependent AAV expressing mCherry into the LS of NTS-Cre mice. We found dense axonal projections of these neurons in the hypothalamus, in particular the lateral hypothalamus (LH, Fig.6A). We also confirmed this projection by injecting a Cre-dependent anterograde herpes simplex virus, H129ΔTK-tdTomato into the LS of NTS-Cre mice (Fig. 6B). A similar LS to LH projection was identified when mapping axonal projections from Glp1r-cre mice expressing a Cre-dependent mCherry from an AAV injected into the LS (Figure S5A). The LH is a well-established site regulating food intake (Morrison et al., 1958), and we then tested whether specific activation of an LS^NTS^ →LH projection would lead to a similar decrease in food consumption as activation of LS^NTS^ cell bodies. We injected a Cre-dependent AAV expressing a channelrhodopsin (Chr2) into the LS of Nts-Cre mice and implanted a fiber above the LH (Figure S5B). Optogenetic activation of this pathway with blue light led to a significant increase in c-fos expression in the LS^NTS^ neurons (∼181 c-fos+ neurons for Chr2 vs. ∼8.5 for mCherry) and a subset of neurons in the LH (∼128 c-fos+ neurons for Chr2 vs. ∼35 for mCherry, Figure S5C and D, Unpaired Students t-test, **p<0.01). We then used an 1-h OFF-ON-OFF protocol to evaluate food intake before, during and after light stimulation of LS^NTS^ terminals in the LH and observed a reversible and significant decrease of food intake in mice expressing ChR2 compared to controls expressing mCherry (Fig. 6C, Two-way ANOVA with post-hoc Bonferroni correction, *p<0.05). Furthermore, similar to the effect of activating the entire LS^NTS^ population of neurons, activation of the LH-projecting LS^NTS^ neurons led to a ∼50% decrease in food intake compared to controls (Fig. 6C, right panel, Unpaired Student’s t-test). We also evaluated whether the activation of LH-projecting LS^NTS^ neurons had any effect on locomotion or anxiety and found no differences during open field or elevated plus maze tests (Figure S5E-G).

**Figure 6.**
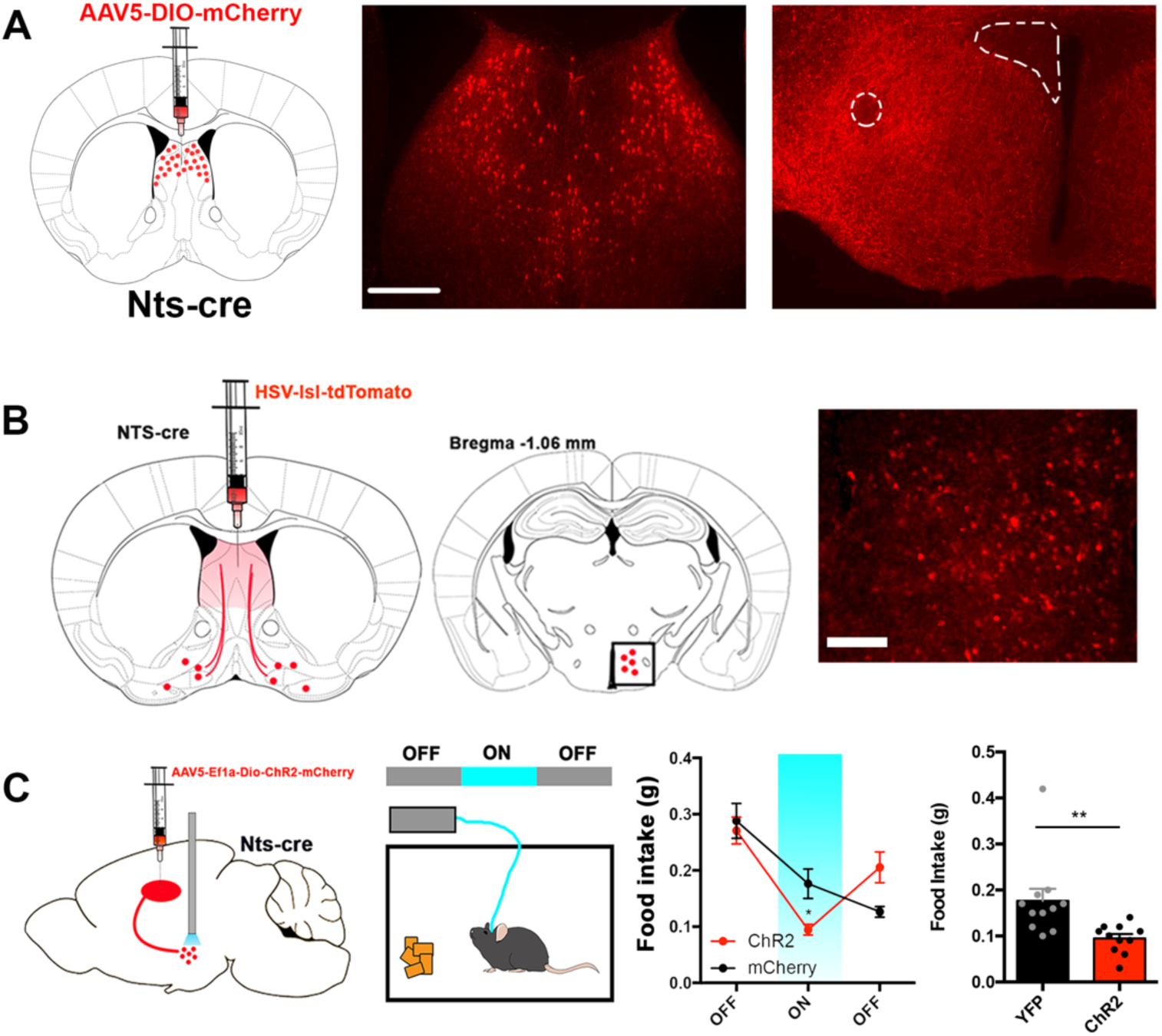
LS^NTS^ ->LH projection regulates food intake in mice. (A) Representative images of the LS injection site (left) and terminals in the lateral hypothalamus (left) of NTS-cre mice injected with a control mCherry reporter virus. Atlas images above the representative images depict the area of magnification. (B) Representative image of cells labeled in the lateral hypothalamus (right) of NTS-cre mice injected with the anterograde tracer HSV-lsl-tdTomato in the LS. Middle, atlas image depicting the area of magnification to the right. (C) Left, experimental scheme depicting mice injected with AAV expressing Chr2 or control mCherry into the LS and an optic fiber positioned over the lateral hypothalamus. Middle, food intake was measured in an Off-On-Off paradigm. Right, food intake measured in grams (g) of control mCherry (black) or Chr2 (red) expressing mice before (off), during (on) and after (off) laser stimulation. n=11, Two-way ANOVA with post-hoc Bonferroni correction, Epoch: F(2,40)=38.47, p<0.0001; Subject: F(1,20)=0.069, p=0.796; Interaction: F(2,40)=10.94, p=0.0002. Data are represented as mean +- SEM.

## DISCUSSION

Stress elicits a pleiotropic set of adaptive physiologic and behavioral responses enabling animals to rapidly respond to threatening situations. These responses include a suppression of other behaviors such as feeding while animals initiate defensive behaviors. Accordingly, the inhibition of consummatory behaviors when animals are threatened decreases an animal’s vulnerability to predation (Petrovich et al., 2009; Kunwar et al., 2015). While acute stress is generally adaptive, chronic stress can increase the risk of maladaptive behaviors and psychiatric disorders including anorexia nervosa. We previously found that neurons in the lateral septum, which is known to mediate the response to stress, can also regulate food intake (Azevedo et al., 2019). We thus considered the possibility that neurons in the LS might link stress to anorexia. We used an unbiased screen to identify molecular markers of neurons that are activated by stress in the LS and identified a novel population of neurons in the LS that co-express of Nts and Glp1r (LS^NTS^ neurons). We further showed that these neurons are activated by active coping in response to stress and that they can suppress food intake and body weight via a projection to the LH. These findings thus identify a molecularly defined neural circuit that responds to stressful stimuli and in turn specifically regulates feeding, but not other components of the stress response.

These data are consistent with a number of prior studies suggesting that the LS plays a role in the active coping response to stress (Singewald et al., 2011; Bondi et al., 2007) though the cell types in the LS that mediate this response had not been identified. In our study, we show a role for LS^NTS^ neurons in this response and also characterized the calcium dynamics of their response using *in vivo* recordings. Consistent with a role in active coping, we found that these neurons are specifically activated only in response to stressful stimuli associated with active movement (i.e; flight) and not with stress associated with freezing/immobility. LS^NTS^ neurons were activated during active escape in a predator-attack paradigm, during tail suspension and during the startle response to a foot shock. In contrast, the immobility associated with complete restraint or freezing after fear conditioning did not lead to activation of these neurons. This finding was initially surprising because our prior results, which mapped c-fos, had shown activation following restraint stress. However, the temporal activation of c-fos is limited and did not allow us to discern whether the LS^NTS^ neurons were activated while the animal was moving or completely immobile. Furthermore, in the initial studies using c-fos as a marker for neural activation, mice were placed in flexible restraint cones which do not completely immobilize them and allows for movement/struggle during the period of restraint. In contrast, in our *in vivo* calcium recording studies, mice were manually restrained under the experimenter’s hand and we observed that mice did not actively struggle during this type of restraint, raising the possibility that manual restraint either does not cause significant stress or that the stress this elicits acts independently of LS^NTS^ neurons.

LS^NTS^ neurons co-express Glp1r are stress-responsive and consistent with our results, a previous report showed that infusion of a Glp1r antagonist (Ex9) directly into the LS diminished the anorexigenic effects of stress, suggesting that Glp signaling in LS contributes to this response (Terril et al., 2018). The LS is innervated by pre-proglucagon (PPG) neurons in the brainstem expressing Glp-1 (Trapp and Cork, 2015) and these neurons have also been shown to be activated by stress (Holt et al., 2019). In aggregate, these data suggest that LS^NTS^ neurons that expresses Glp1r provide a link between brainstem circuits conveying stress to key output pathways from brainstem and perhaps elsewhere to regulate feeding. Further studies mapping the inputs to LS^NTS^ neurons will be necessary to confirm this possibility.

LS^NTS^ neuronal activation decreases food intake suggesting that cells may be tuned by active coping mechanisms that dispense with voluntary food intake until appropriate countermeasures to mitigate the stressful event have been enacted. For example, this circuit would be useful for suppressing the urge to forage when predators are in the vicinity, thus serving an important evolutionary function. Consistent with this, activation of LS^NTS^ neurons suppresses food intake even in animals that have been food deprived. If true, neural plasticity and/or genetic alterations within this circuit (e.g. by mutations in the Nts gene [Lutter et al., 2017] or changes in “allostatic load” (McEwen and Akil, 2020) could contribute to persistent maladaptive eating strategies, such as those observed in eating disorders or in other cases obesity. Presumably other neural circuits mediate the effects of stressful stimuli that lead to freezing raising the possibility that different circuits have evolved to process different types of stress, and that different types of chronic stress might be associated with different disorders.

Indeed, a strong association between stress and reduced food intake has been noted in patients with eating disorders, such as Anorexia Nervosa, which has the highest mortality rate of any psychiatric disorder and is more prevalent in females (Treasure et al., 2020; Kaye, 2008). Moreover, Anorexia Nervosa typically manifests during puberty, a critical period characterized by increased levels of estrogen (Kaye, 2008; Ma et al., 2019). Consistent with an effect of sex hormones on this circuit, we found that the activity of LS^NTS^ neurons is increased in stressed females during estrus compared to stressed females in diestrus or compared to males. This suggest that Nts expression in the LS is tightly regulated by estrogen and that both stress and high levels of estrogen are necessary for maximal neural activation. While Nts expression has previously been shown to be regulated by estrogen in the preoptic area of the hypothalamus, a brain area responsible for maternal behavior, Nts neurons in this region have not been shown to regulate feeding (Alexander et al., 1994).

Additional evidence suggesting a possible link between LS^NTS^ neurons and anorexia nervosa has recently been provided in recent WGS (whole genome sequencing) and GWAS (gene-wide association studies) studies of subset of female patients with AN who report restrictive eating behaviors (Lutter et al., 2017; Watson et al., 2019). In particular, one WGS study (Lutter et al., 2017) reported linkage of polymorphisms in Glp1r and Nts to AN. In another GWAS studies (Pinheiro et al., 2010), seven genes were identified that we also find as marking LS neurons that are activated by stress (see Table S1 and Table S2) including *Comt, Rgs10, Pomc, Crh, Calm1, Gnas, Esrra*. Lastly, the most recent GWAS study for anorexia nervosa (Pinheiro et al., 2010) identified 8 significant loci and 4 genes of particular interest. Of these, 2 genes (*Cadm1* and *Mgmt*) and another that was less strongly implicated (*Gpr27*) were significantly increased in Nts neurons according to our transcriptomic data (Table S1) and one (*Foxp1*) was significantly decreased (Table S1). While it is not yet known whether these additional genes can also regulate feeding behavior, these data add further evidence linking the activity of LS^NTS^ neurons to anorexia and possibly eating disorders.

If as suggested by these results, LS^NTS^ neurons play a role in the pathogenesis of anorexia nervosa, reducing their activity pharmacologically could be of therapeutic benefit. Thus, it is possible that pharmacologically targeting Glp-1 signaling in patients with eating disorders, or other means for regulating the activity of LS^NTS^ neurons, could provide important clinical insights and possibly even therapeutic effects. Further studies will be needed to assess this possibility, including analyses of the effect of modulating neurons in animal models of anorexia nervosa, though it is unclear to what extent rodent models of this disorder fully recapitulates the features of the human condition. It has also been suggested that projections from the limbic system to the hypothalamus may mediate the psychological aspects of body weight preoccupation and reduction of feeding in AN (Donohoe et al., 1994), but the identity of these potential afferents had not been characterized. Our finding that LS^NTS^ neurons reduce food intake via projections to the LH represents another potential link to eating disorders. Numerous other studies have also established connections between the lateral septum and the lateral hypothalamus to control food intake (Azevedo et al., 2019; Sweeney and Yang, 2016) as well as other behaviors including cocaine-induced locomotion and reward (McGlinchey et al., 2016; Reddy et al., 2016).

In summary, these data suggest that LS^NTS^ neurons play an important integratory function linking stress to anorexia and serving as an anatomical bridge between the limbic system and the hypothalamus. The identification of these neurons thus provides an entry point for understanding how complex inputs - in this case food deprivation vs. danger - compete to generate an appropriate adaptive response, in some cases potentially leading to a pathologic response in the case of eating disorders. These and other data further (Azevedo et al., 2019; Newmeyer et al., 2019; Land et al., 2014) suggest that higher-order, limbic brain regions integrate diverse sensory information to modulate subcortical feeding circuits in health and possibly disease.

## MATERIALS AND METHODS

### Animals and breeding

C57BL/6J (Stock# 000664), NTS-Cre (Stock# 017525), Sst-Cre (Stock# 028864), Cartpt-Cre (Stock# 028533), and Glp1r-cre (Stock# 029283) mice were obtained from Jackson Laboratories and housed according to the guidelines of the Rockefeller’s University Animal Facility. Males and females at age 8-20 weeks were used throughout this study and all animal experiments were approved by the Rockefeller University IACUC, according to NIH guidelines. Male and female mice were used for NTS-cre, Sst-cre, Cartpt-Cre and Glp1r-cre experiments. In experiments that used Cre+ and Cre-mice, littermates were used. No differences were determined based on gender. Mice were kept on a 12-h/12-h light/dark cycle (lights on at 7:00 a.m.) and had access to food and water ad libitum, except when noted otherwise. All feeding experiments used standard rodent chow pellets.

### Stereotaxic Injections

All AAVs used in this study were purchased from UNC Vector Core or Addgene, except where noted. AAV5-DIO-hM3Dq-mCherry or AAV5-DIO-Chr2-mCherry were used for activation studies. AAV5-DIO-hM4Di-mCherry was used for inhibition studies. Control AAV5-DIO-mCherry virus was used for comparison and for terminal projection mapping. AAV5-DIO-GCaMP6s was used for fiber photometry studies. AAV5-Introvert-GFPL10a was a gift from Alexander Nectow. H129ΔTK-TT virus was a gift from David Anderson and was used as described previously (Lo and Anderson, 2011) Mice were anesthetized with isoflurane, placed in a stereotaxic frame (Kopft Instruments) and bilaterally injected in the lateral septum using the following coordinates relative to bregma: AP: +0.1; ML: 0.0, DV: -3.00 (Paxinos). A total of 400 nL of virus at high titer concentrations (at least 10^11^) were injected per site at a rate of 100 nL/min. For pharmacological experiments, cannulae were implanted in the LS at the same coordinates as above. For fiber photometry experiments, optic fiber implants (Doric) were inserted at the same coordinates as above. For optogenetics experiments, optic fiber implants (Thor Labs) were implanted bilaterally over the LH at the following coordinates relative to bregma: AP: -1.80 mm; ML: + 1.50 mm, DV: -5.25 mm at a 10^°^ angle (Paxinos). Implants were inserted slowly and secured to the mouse skull using two layers of Metabond (Parkell Inc). Mice were singly housed and monitored in the first weeks following optic fiber implantation. For all surgeries, mice were monitored for 72h to ensure full recovery and two weeks later mice were used in experiments. Following behavioral procedures, mice were euthanized to confirm viral expression and fiber placement using immunohistochemistry.

### Restraint Stress

Except where noted for fiber photometry studies, restraint stress consisted of 1 hour of immobilization in a decapicone (Braintree Scientific) in a separate room adjacent to the housing room. Mice were returned to their home cage following restraint for further food intake measurements or until euthanasia. For in vivo fiber photometry, restrain stress was given manually by the experimenter during recordings.

### Fiber photometry

Mice were acclimated to patch cables for 5 min for 3 days before experiments. Analysis of the signal was done using fiber photometry system and processor (RZ5P) from TDT (Tucker-Davis Technologies), which includes the Synapse software (TDT). Post-recording analysis was performed using custom-written MATLAB codes. The bulk fluorescent signals from each channel were normalized to compare across animals and experimental sessions. The 405 channel was used as the control channel. GCaMP6s signals that are recorded at this wavelength are not calcium-dependent; thus, changes in signal can be attributed to autofluorescence, bleaching, and fiber bending. Accordingly, any fluctuations that occurred in the 405 control channels were removed from the 465 channel before analysis. Change in fluorescence (ΔF) was calculated as (465 nm signal - fitted 405 nm signal), adjusted so that a ΔF/F was calculated by dividing each point in ΔF by the 405-nm curve at that time. Z-score, plot traces and colormaps were calculated for all experiments using MATLAB. Behavioral variables were timestamped in the signaling traces via the real-time processor as TTL signals from Noldus (Ethovision) software. This allowed for precise determinations of the temporal profile of signals in relation to specific behaviors. Behaviors were tested as follows. First, we tested response to food. Mice were fasted overnight and then placed into a clean home cage with food pellets introduced 5 minutes later. Food approach and consumption were scored. We then tested restraint by manually restraining mice for 5 minutes. Tail suspension was conducted by suspending mice by their tails for 5 minutes. To examine the role of predator escape, mice were placed into an open field that contained a designated safe zone consisting of an acrylic “nest”. After 5 minutes, a remote-controlled spider (Lattburg) was introduced into the open field and began to pursue the mouse at random intervals. After 10-15 minutes the nest was removed. We scored times when the mouse fled from the spider as well as the removal of the nest. Locomotion was scored during the first 5 minutes in the open field arena. All behavior was manually scored by a second investigator who did not conduct the photometry experiment.

### Chemogenetic and Pharmacological Feeding Experiments

For acute feeding, mice were single housed and injected with either saline or clozapine-N-oxide (CNO; Sigma) at 1 mg/kg intraperitoneally (i.p), exendin-4 (Tocris, 1 mg/mL), or y-msh (Tocris, 150 nM) directly into the lateral septum before the dark phase and cumulative food intake and body weight was measured at selected time periods. For chronic feeding, mice were single or double injected (every 12h) with CNO or saline before dark phase and food intake and body weight were measured. For experiments that require fasting, mice were single housed and food was removed before dark phase (16-23h of fasting in total). Cumulative food intake at each time point was calculated for analysis.

### Elevated Plus Maze

Mice were placed at the center of a cross-shaped, elevated maze in which two arms are closed with dark walls and two arms are open and allowed to explore for 10 min. Mice were injected with CNO (1mg/kg) 1h before testing. After, mice were returned to their home cage and the maze floor was cleaned in between subjects. All subjects were recorded using a camera and behavior (time spent in open and closed arms, distance and velocity) were analyzed using Ethovison 9.0 (Noldus).

### Open Field / Open Field with Novelty

Mice freely explored an open field arena (28 x 28 cm), divided into center and border regions, for 5 min. The time spent (in seconds) in the center area, were taken as measures of anxiety. To increase the anxiolytic nature of the task, a novel object was placed into the center of the arena (open field with novelty). Mice were injected with CNO (1mg/kg) 1h before testing. After, mice were returned to their home cage and the maze floor was cleaned in between subjects. All subjects were recorded using a camera and behaviors (time spent in center and border, distance and velocity) were analyzed using Ethovison 9.0 (Noldus).

### Optogenetics Feeding Experiment

Mice were previously habituated to patch cables for 3 days before experiments. Implanted optic fibers were attached to a patch cable using ceramic sleeves (Thorlabs) and connected to 473 nm laser (OEM Lasers/OptoEngine). Laser output was verified at start of each experiment. Blue light was generated by a 473 nm laser diode (OEM Lasers/OptoEngine) at 5-10 mW of power. Light pulse (10 ms) and frequency (10Hz) was controlled using a waveform generator (Keysight) to activate NT+ terminals in lateral hypothalamus. Animals were sacrificed to confirm viral expression and fiber placement using immunohistochemistry. Each feeding session lasted 80 min and it was divided in 1 trial of 20 min to allow the animal to acclimate to the cage and 3 trials of 20 min each (1h feeding session). During each feeding session, light was off during the first 20 min, on for 20 min and off again for the remaining 20 min. Consumed food was recorded manually before and after each session. To facilitate measurement, 3 whole pellets were added to cups and food crumbs were not recorded. Feeding bouts were recorded using a camera and the Ethovision 9.0 software (Noldus).

### Viral-Trap (V-Trap)

Affinity purification of EGFP-tagged polysomes was done 3 or 4 weeks after virus injections (Nectow et al., 2017). Briefly, mice were separated into 3 biological replicate groups of 4-6 mice per group and euthanized. Brains were removed, and the lateral septum was dissected on ice and pooled. Tissue was homogenized in buffer containing 10 mM HEPES-KOH (pH 7.4), 150 mM KCl, 5 mM MgCl2, 0.5 mM DTT, 100 μg/mL cycloheximide, RNasin (Promega) and SUPERase-In™ (Life Technologies) RNase inhibitors, and Complete-EDTA-free protease inhibitors (Roche) and then cleared by two-step centrifugation to isolate polysome-containing cytoplasmic supernatant. Polysomes were immunoprecipitated using monoclonal anti-EGFP antibodies (clones 19C8 and 19F7; see Heiman et al., 2008^30^) bound to biotinylated-Protein L (Pierce; Thermo Fisher Scientific)-coated streptavidin-conjugated magnetic beads (Life Technologies). A small amount of tissue RNA was saved before the immunoprecipitation (Input) and both input and immunoprecipitated RNA (IP) were then purified using RNAeasy Mini kit (QIAGEN). RNA quality was checked using an RNA PicoChip on a bioanalyzer. RIN values >7 were used. Experiments were performed in triplicates for each group. cDNA was amplified using SMARTer Ultralow Input RNA for Illumina Sequencing Kit and sequenced on an Illumina HiSeq2500 platform.

### RNA sequencing and qPCR analysis

RNA sequencing raw data was uploaded and analyzed using BaseSpace apps (TopHat and Cufflinks; Illumina) using an alignment to annotated mRNAs in the mouse genome (UCSC, Mus musculus assembly mm10). The average immunoprecipitated (IP) and Input value of each enriched and depleted genes with a q-value lower than 0.05 were plotted using GraphPad Prism (GraphPad). To validate the RNA Sequencing data, qPCR using predesigned Taqman probes (idtDNA) were used and cDNA was prepared using the QuantiTect Reverse Transcription Kit (Life Technologies). The abundance of these genes in IP and Input RNA was quantified using Taqman Gene Expression Master Mix (Applied Biosystems). Transcript abundance was normalized to beta-actin. Fold of Change were calculated using the ΔΔCt method.

### Immunohistochemistry, quantifications and imaging

Mice were perfused, and brains were postfixed for 24h in 10% formalin. Brain slices were taken using a vibratome (Leica), blocked for 1h with 0.3% Triton X-100, 3% bovine serum albumin (BSA), and 2% normal goat serum (NGS) and incubated in primary antibodies for 24h at 4°C. Then, free-floating slices were washed three times for 10 min in 0.1% Triton X-100 in PBS (PBS-T), incubated for 1h at room temperature with secondary antibodies, washed in PBS-T and mounted in Vectamount with DAPI (Southern Biotech). Antibodies used here were: anti-c-fos (1:500; Cell Signaling), anti-mCherry (1:1000; Abcam) goat-anti-rabbit (Alexa 488 or Alexa 594, 1:1000; Molecular Probes), goat anti-chicken Alexa488 or Alexa594 (1:1000; Molecular Probes). Images were taken using Axiovert 200 microscope (Zeiss) or LSM780 confocal (Zeiss) and images were processed using ImageJ software (NIH, Schneider et al., 2012). C-fos counts were conducted for 3 or more sections / animal and averaged for each animal for statistical analysis.

### Fluorescent In Situ Hybridization

For examination of gene expression and Fos experiments, tissue samples underwent single molecule fluorescent in situ hybridization (smFISH). Isoflurane anesthetized mice were decapitated, brain harvested and flash frozen on aluminum foil on dry ice. Brains were stored at -80°C. Prior to sectioning, brains were equilibrated to -16°C in a cryostat for 30 min. Brains were cryostat sectioned coronally at 20 μm and thaw-mounted onto Superfrost Plus slides (25×75 mm, Fisherbrand). Slides were air-dried for 60 to 90 min prior to storage at -80°C. smFISH for all genes examined— *Fos, Nts, Sst, Glp1r, Cartpt, Mc3r* - was performed using RNAscope Fluorescent Multiplex Kit (Advanced Cell Diagnostics) according to the manufacturer’. Slides were counterstained for the nuclear marker DAPI using ProLong Diamond Antifade mounting medium with DAPI (Thermo Fisher). Sections were imaged using an LSM780 confocal (Zeiss) and processed using ImageJ software. Counts were conducted for 2 or more sections/animal and averaged for each animal for statistical analysis.

### Statistical analysis

All results are presented as mean ± SEM. and were analyzed with Prism software or with Matlab. No statistical methods were used to predetermine sample sizes, but our sample sizes are similar to those reported in previous publications. Normality tests and F tests for equality of variance were performed before choosing the statistical test. Unless otherwise indicated, for feeding experiments we used a two-way ANOVA with Bonferroni correction to analyze statistical differences. P < 0.05 was considered significant (*p < 0.05, **p< 0.01, ***P < 0.001, ****p<0.0001). Image quantifications were analyzed using unpaired Students’ t-tests. Fiber photometry data were analyzed using paired Students’ t-tests. Animals in the same litter were randomly assigned to different treatment groups and blinded to experimenters in the various experiments. Injection sites and viral expression were confirmed for all animals. Mice showing incorrect injection sites or optic fiber placement were excluded from the data analysis.

## ACKNOWLEDGEMENTS

We thank Ravi Tolwani and the staff of the Comparative Biosciences Center, Connie Zhao and the staff of the Genomics Resource Center, Alison North and the staff of the Bioimaging Resource Center at Rockefeller University for technical assistance. We thank Alexander Nectow for the AAV-Introvert-GFPL10a virus and David Anderson for providing the HSV-lsl-tdTomato virus. We thank Christin Kosse for providing MATLAB codes for photometry analysis. This work was funded by a Long Term HSFP Fellowship (E.P.A.), F32DK107077 (S.A.S.), NARSAD Young Investigator Awards from the Brain and Behavior Research Foundation (E.P.A and S.A.S) the JPB Foundation and the Klarman Foundation (J.M.F).

## SUPPLEMENTARY FIGURES

**Supplementary Figure S1.**
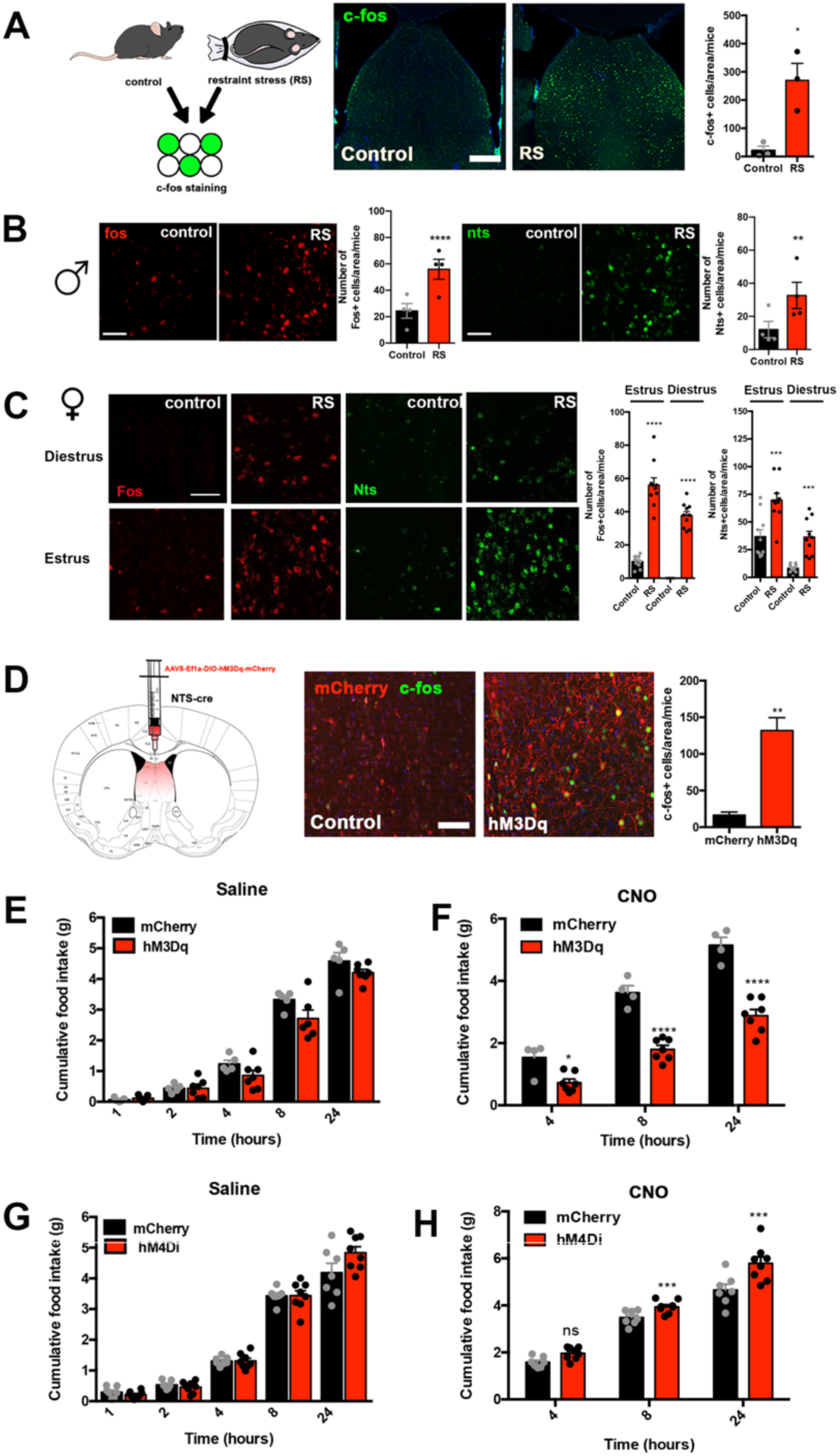
LS^NTS^ neurons regulate feeding behavior. (A) Representative images of c-fos (green) in the lateral septum of control and restraint stress (RS) mice. Right, quantification of c-fos+ cells in control (black) and RS (red) mice. n= 4, Unpaired Student’s t-test, *p<0.05, t=3.964, df=4. Scale bars, 50 um. (B) Representative images of in situ hybridization experiments measuring the levels of fos (red) and Nts (green) in the lateral septum of control and restraint stress (RS) male mice. Left, quantification of fos+ cells in control (black) and RS (red) male mice, n= 4, Nested t-test, ****p<0.0001, t=5.175, df=34. Right, quantification of nts+ cells in control (black) and RS (red) male mice, n= 4, Nested t-test, **p<0.01, t=3.476, df=34. Scale bars, 50 um. (C) Representative images of in situ hybridization experiments measuring the levels of fos (red) and Nts (green) in the lateral septum of control and restraint stress (RS) female mice in estrus or diestrus phase. Left, quantification of fos+ cells in control (black) and RS (red) female mice in estrus or diestrus phase. n= 4, ****p<0.0001, F(3,34)=90.19. Right, quantification of nts+ cells in control (black) and RS (red) female mice in estrus or diestrus phase. n= 4, ****p<0.0001, F(3,38)=28.18. Scale bars, 50 um. (D) Representative images of viral injection strategy and viral expression (mCherry, red) and c-fos (green) in control virus (left) and hM3Dq (right) injected mice following CNO injection. Right, quantification of c-fos+ cells in mCherry control (black) and hM3Dq (red) expressing mice. n=3, Unpaired Students’ t-test, p=0.003, t(4)=6.418. (E) Food intake, measured in grams (g), in control mCherry (black) and hM3Dq (red) expressing Nts-cre mice 1 hour (h), 2h, 4h, 8h and 24h following saline injection. n=5(mCherry, n=5(hM3Dq). Mixed-model ANOVA with post-hoc Bonferroni correction (Time: F(2.33,22.72)=298.8, p<0.0001; Subject: F(1,10)=6.203, p=0.032; Interaction: F(4,39)=1.727, p=0.1637. (F) Food intake, measured in grams (g), in control mCherry (black) and hM3Dq (red) expressing Nts-cre mice 4 hour (h), 8h and 24h following CNO injection. n=7(mCherry), n=10(hM3Dq). Two-way ANOVA with post-hoc Bonferroni correction (Time: F(2,18)=, p<0.0001; Subject: F(1,9)=60.03, p<0.0001; Interaction: F(2,18)=12.38, p=0.0004. (G) Food intake, measured in grams (g), in control mCherry (black) and hM4Di (red) expressing Nts-cre mice 1 hour (h), 2h, 4h, 8h and 24h following saline injection. n=7(mCherry), n=8(hM4Di). Two-way ANOVA with post-hoc Bonferroni correction (Time: F(1,19)=415.8, p<0.0001; Subject: F(1,13)=1.042, p=0.326; Interaction: F(4,52)=2.839, p=0.0333. (H) Food intake, measured in grams (g), in control mCherry (black) and hM4Di (red) expressing Nts-cre mice 4 hours (h), 8h and 24h following CNO injection. n=7(mCherry), n=8(hM3Dq). Two-way ANOVA with post-hoc Bonferroni correction (Time: F(2,18)=184.5, p<0.0001; Subject: F(9,18)=2.945, p=0.0245; Interaction: F(2,18)=12.38, p=0.004.

**Supplementary Figure S2.**
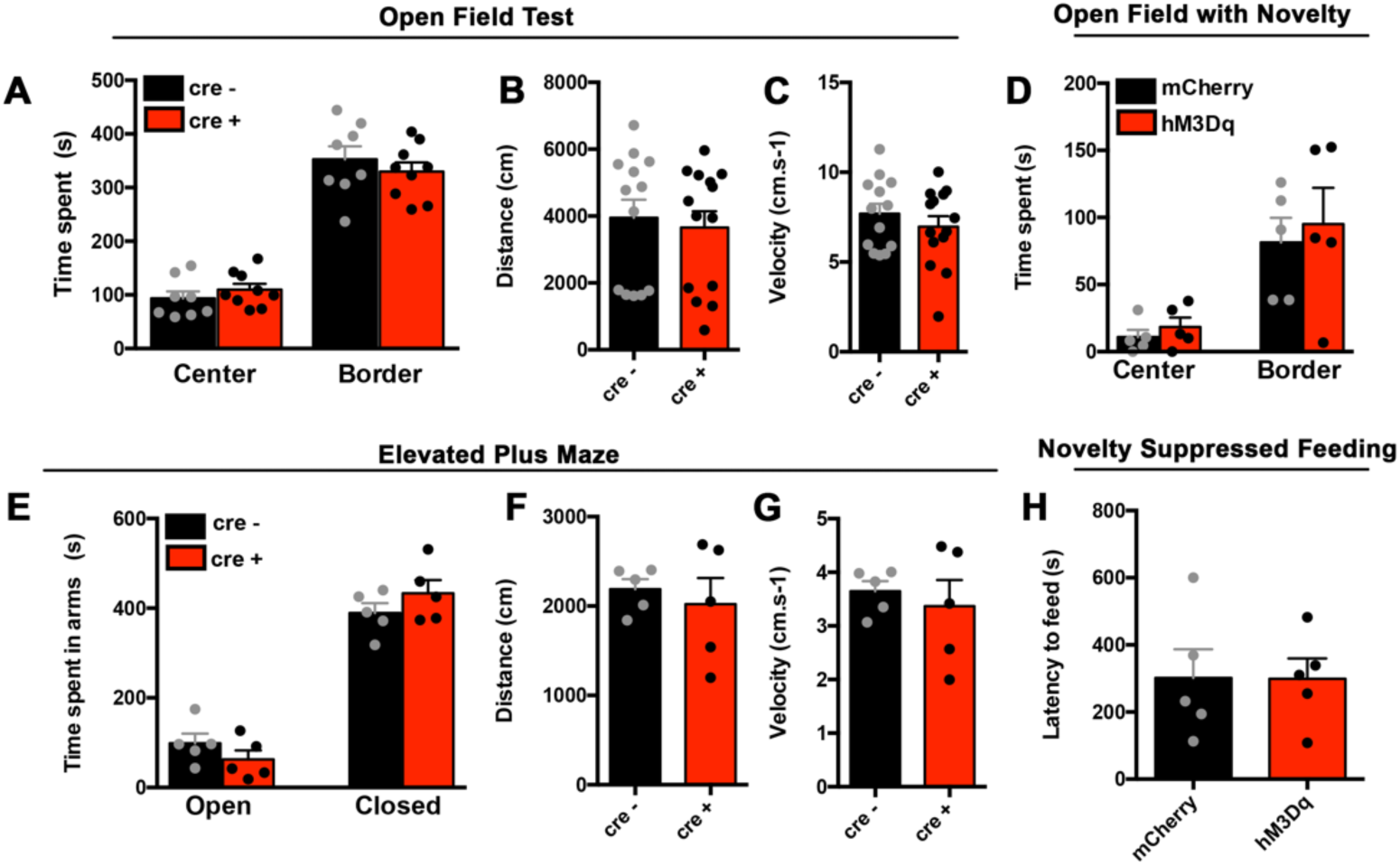
LS^NTS^ neurons do not regulate anxiety-like behaviors. (A) Time, measured in seconds (s), spent in the center and border of an open field in control mCherry (black) and hM3Dq (red) expressing Nts-cre mice. n=8(mCherry), n=9(hM3Dq). Two-way ANOVA with post-hoc Bonferroni correction, Subject: F(1,15)=0.071, p=0.794; Arena: F(1,15)=135.8, p<0.0001; Interaction: F(1.15)=0.9093, p=0.355. (B) Distance, measured in centimeters (cm), traveled in the open field, in control mCherry (black) and hM3Dq (red) expressing Nts-cre mice. n=13(mCherry), n=14(hM3Dq), unpaired Students’ t-test, t(25)=0.403, p=0.69. (C) Velocity, measured as the cm per s (cm.s^-1^), in the open field, in control mCherry (black) and hM3Dq (red) expressing Nts-cre mice. n=13(mCherry), n=14(hM3Dq), unpaired Students’ t-test, t(25)=0.915, p=0.369. (D) Time, measured in seconds (s), spent in the center and border of an open field with novelty in control mCherry (black) and hM3Dq (red) expressing Nts-cre mice. n=5. Two-way ANOVA with post-hoc Bonferroni correction, (Subject: F(1,8)=0.358, p=0.566; Arena: F(1,8)=21.54, p=0.0017; Interaction: F(1,8)=0.039, p=0.847. (E) Time, measured in seconds (s), spent in the open and closed arms of an elevated plus maze in control mCherry (black) and hM3Dq (red) expressing Nts-cre mice. n=5. Two-way ANOVA with post-hoc Bonferroni correction, Arm: F(1,8)=111.2, p<0.0001; Subject: F(1,8)=0.137, p=0.72; Interaction: F(1,8)=1.641, p=0.236 (F) Distance, measured in centimeters (cm), traveled in the elevated plus maze, in control mCherry (black) and hM3Dq (red) expressing Nts-cre mice. n=5, unpaired Students’ t-test, t(8)=0.5347, p=0.6074. (G) Velocity, measured as the cm per s (cm.s^-1^), in the elevated plus maze, in control mCherry (black) and hM3Dq (red) expressing Nts-cre mice. n=5, unpaired Students’ t-test, t(8)=0.533, p=0.608. (H) Latency to feed, measured in seconds (s), in a novelty suppressed feeding task, in control mCherry (black) and hM3Dq (red) expressing Nts-cre mice. n=5, unpaired Students’ t-test, t(8)=0.027, p=0.979. Data are represented as mean +- SEM.

**Supplementary Figure S3.**
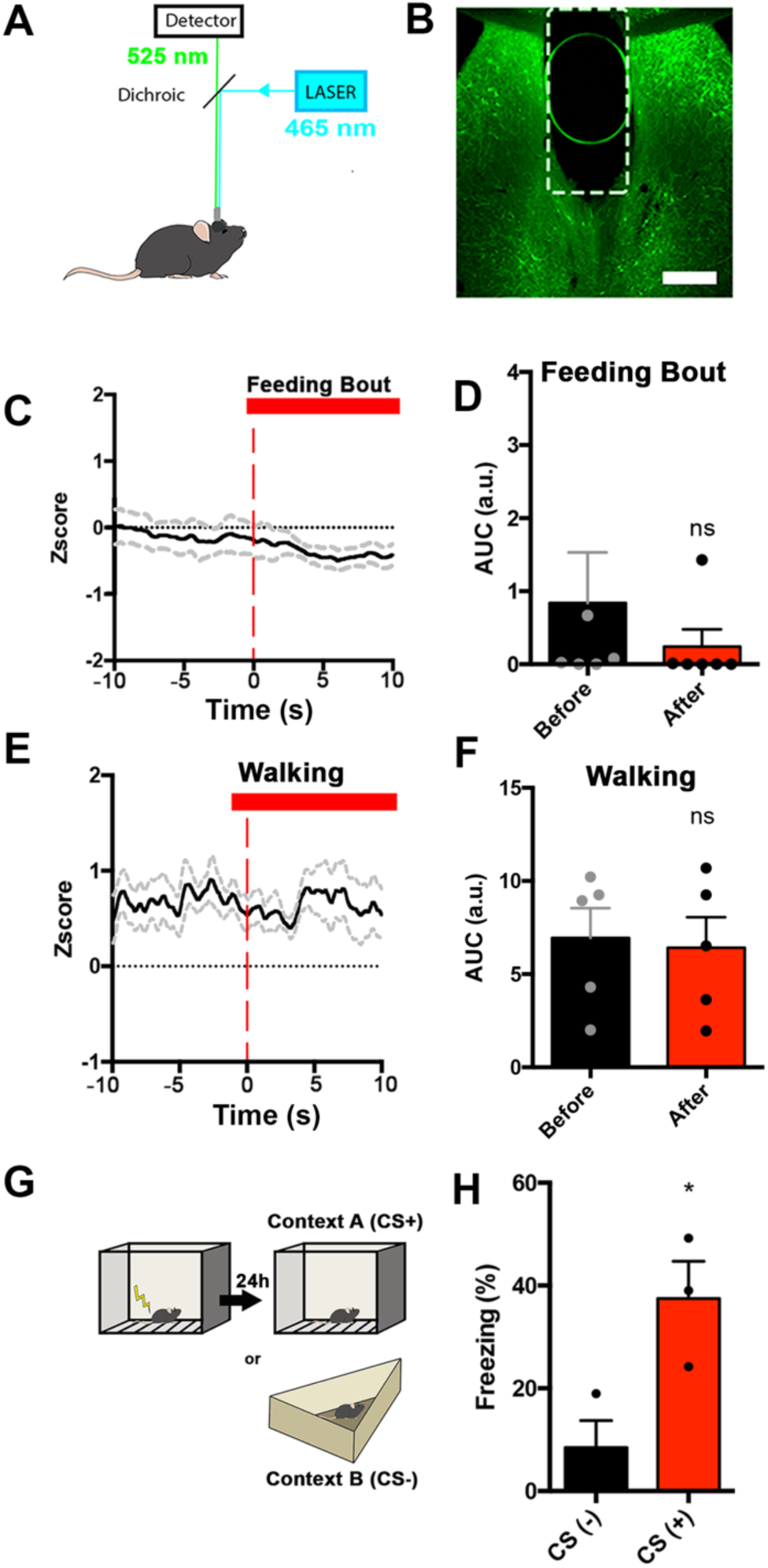
LS^NTS^ neurons are not selectively tune to feeding or movement. (A) Experimental scheme. Mice were then attached to a fiber photometry setup with a 465nm LED (blue) and a 405 nm LED (purple) and a photodetector. (B) Representative image of viral expression in the LS of Nts-cre mice injected with gCaMP6s and implanted with an optic fiber in the LS. (C) Average gCaMP6s z-score (black lines) and SEM (gray dotted lines) across all recording sites time-locked to the start of the feeding bout. Right, Heat maps represent the average % DF/F from each gCaMP6s recordings time-locked to feeding bout start. (D) Area under the curve (a.u.) of fluorescent signal before (black) and after (red) the feeding bout start. n=6, Paired Students’ t-test, t(5)=1.304, p=0.249, (E) Average gCaMP6s z-score (black lines) and SEM (gray dotted lines) across all recording sites time-locked to the start of the start of locomotion. (F) Area under the curve (a.u.) of fluorescent signal before (black) and after (red) the start of locomotion. n=5, Paired Students’ t-test, t(4)=0.402, p=0.7083. (G) Experimental scheme of contextual fear conditioning. (H) Percentage of freezing in Context A (CS+, red) and Context B (CS-, black). Unpaired Student’s t-test, *p<0.05, t=3.233, df=4, n=3. Data are represented as mean +- SEM.

**Supplementary Figure S4.**
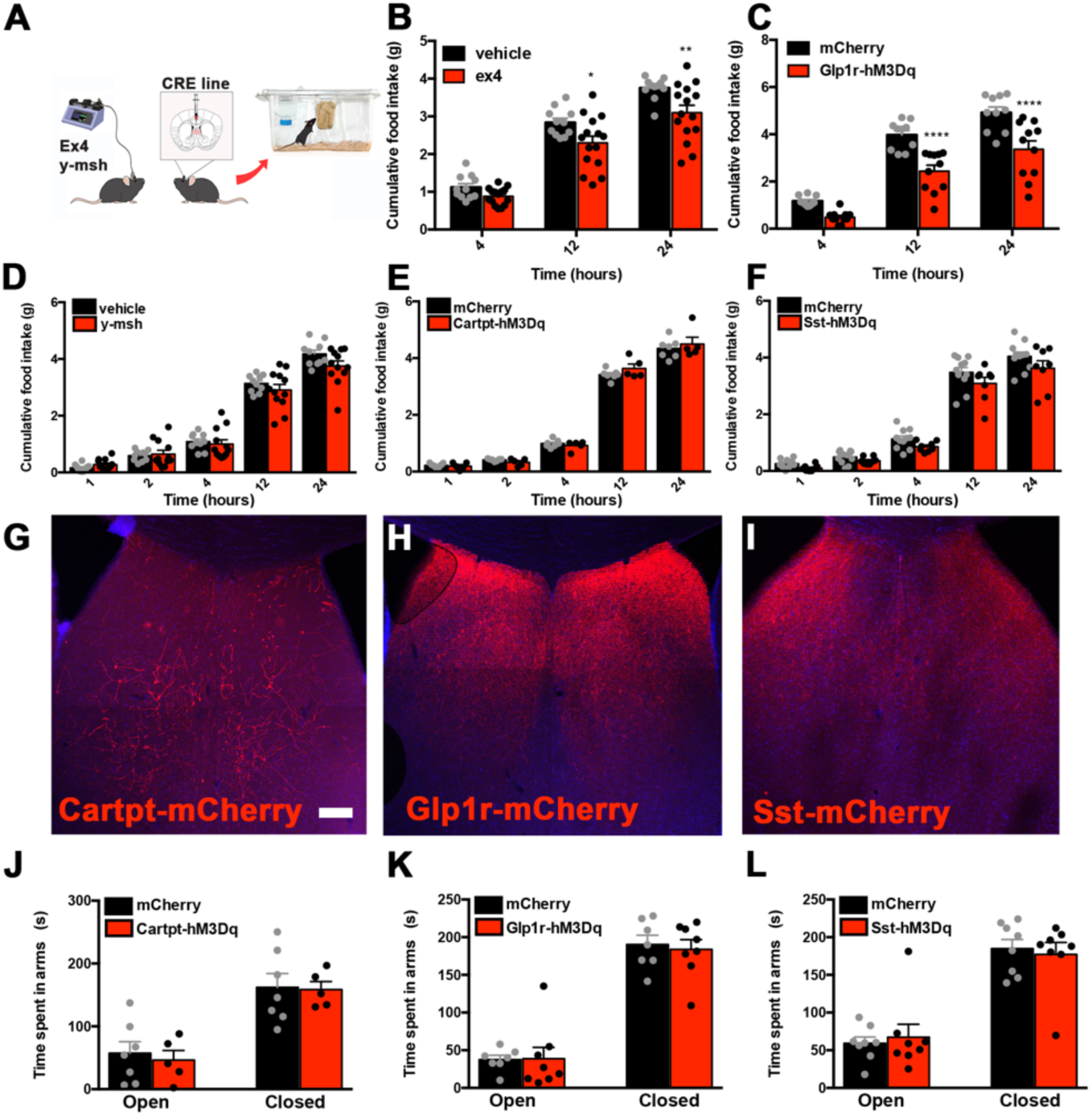
Regulation of feeding and/or anxiety by Cartpt-, Glp1r- and Sst-expressing neurons in the lateral septum. (A) Experimental scheme. Wild-type mice are injected with Ex4 or Mc3r agonist y-MSH directly into the LS or Glp1r-cre, Cartpt-Cre or Sst-cre mice with viral expression of hM3Dq activatory DREADD receptor are tested for cumulative food intake. (B) Food intake, measured in grams (g) of mice 4 hours (h), 12h and 24h following Vehicle (black) and Ex4 (red) injection in the LS. n=11(Vehicle), n=15(Control), Two-way ANOVA with post-hoc Bonferroni correction (Time: F(2,48)=223.6, p<0.0001; Treatment: F(1,24)=9.904, p=0.004; Interaction: F(2,48)=1.67; p=0.199. (C) Food intake, measured in grams (g) of control mCherry (black) and hM3Dq (red) expressing Glp1r-cre mice 4 hours (h), 12h and 24h following CNO injection. n=xx, Two-way ANOVA with post-hoc Bonferroni correction. n=10(mCherry), n=11(hM3Dq). (Time: F(2,38)=296.4, p<0.0001; Subject: F(1,19)=20.32, p=0.0002; Interaction: F(2,38)=6.077, p=0.0051. (D) Food intake, measured in grams (g) of mice 1-hour (h), 2h, 4h, 12h and 24h following Vehicle (black) and y-msh (red) injection in the LS. n=10(Vehicle), n=12(y-msh), Two-way ANOVA with post-hoc Bonferroni correction (Time: F(4,80)=585.6, p<0.0001; Treatment: F(1,20)=0.66, p=0.426; Interaction: F(4,80)=2.67; p=0.038. (E) Food intake, measured in grams (g) of control mCherry (black) and hM3Dq (red) expressing Cartpt-cre mice 1 hour (h), 2h, 4h, 12h and 24h following CNO injection n=7(mCherry), n=5(hM3Dq), Two-way ANOVA with post-hoc Bonferroni correction. (Time: F(2,17)=1,075, p<0.0001; Subject: F(1,10)=0.1767, p=0.6831; Interaction: F(4,40)=1.364, p=0.2637. (F) Food intake, measured in grams (g) of control mCherry (black) and hM3Dq (red) expressing Sst-cre mice 1 hour (h), 2h, 4h, 12h and 24h following CNO injection. n=xx, Two-way ANOVA with post-hoc Bonferroni correction. n=11(mCherry), n=12(hM3Dq). (Time: F(4,68)=660.3, p<0.0001; Subject: F(1,17)=3,23, p=0.0903; Interaction: F(4,68)=1.006, p=0.4106. (G) Representative image of viral expression in the lateral septum of Cartpt-cre mice. (H) Representative image of viral expression in the lateral septum of Glp1r-cre mice. (I) Representative image of viral expression in the lateral septum of Sst-cre mice. (J) Time, measured in seconds (s), spent in the open arms and closed arms of an elevated plus maze in control mCherry (black) and hM3Dq (red) expressing Cartpt-cre mice.n=7(mCherry), n=5(hM3Dq), Two-way ANOVA with post-hoc Bonferroni correction. (Arm: F(1,10)=17.12, P<0.0001; Subject: F(1,10)=1.621, P=0.232; Interaction: F(1,10)=0.02, p=0.891. (K) Time, measured in seconds (s), spent in the open arms and closed arms of an elevated plus maze in control mCherry (black) and hM3Dq (red) expressing Glp1r-cre mice. n=7(mCherry), n=8(hM3Dq), Two-way ANOVA with post-hoc Bonferroni correction. (Arm: F(1,13)=79.20, P<0.0001; Subject: F(1,13)=0.3198, P=0.5814; Interaction: F(1,13)=0.05114, p=0.825. (L) Time, measured in seconds (s), spent in the open arms and closed arms of an elevated plus maze in control mCherry (black) and hM3Dq (red) expressing Sst-cre mice. n=8, Two-way ANOVA with post-hoc Bonferroni correction. (Arm: F(1,14)=39.48, p<0.0001; Subject: F(1,14)=0.0007, p=0.979; Interaction: F(1,14)=0.1811, p=0.677. Data are represented as mean +- SEM.

**Supplementary Figure S5.**
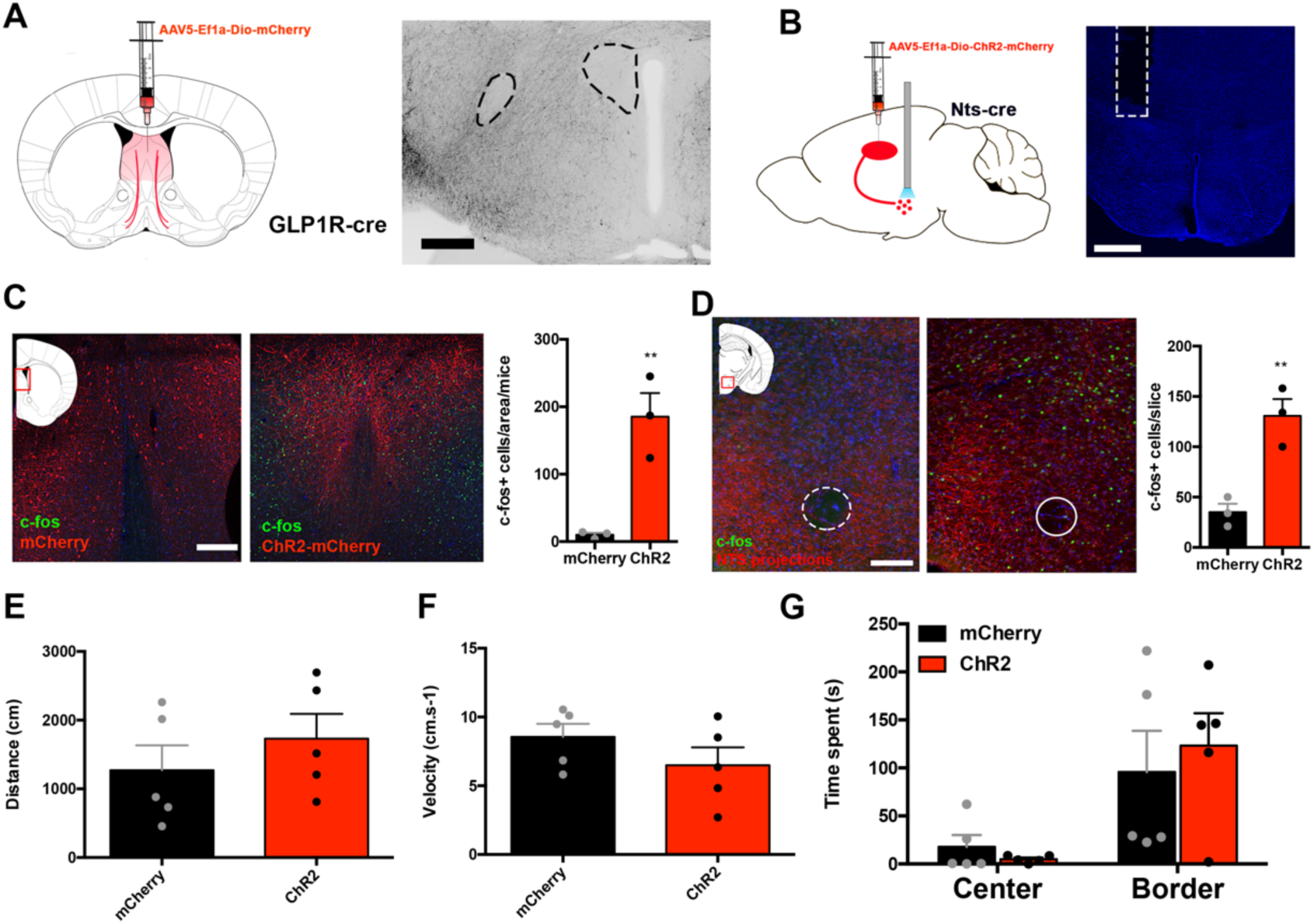
LS^NTS^ → LH does not regulate anxiety or locomotion. (A) Left, experimental scheme of Glp1r-cre mice injected with AAV expressing cre-dependent mCherry in the lateral septum. Right, representative image of Glp1r fibers in the lateral hypothalamus. Scale bar are 100 um. (B) Left, experimental scheme of Nts-cre mice injected with AAV expressing Chr2 in the lateral septum and optic fibers in the lateral hypothalamus. Right, Representative image of fiber optic path in the lateral hypothalamus. Scale bar are 100 um. (C) Left, representative images of c-fos (green) and mCherry (red) in control mCherry (left) and Chr2 (right) expressing Nts-cre mice injected with CNO. Right, quantification of c-fos+ cells in control (black) and Chr2 (red) mice. n=3, unpaired Students’ t-test, t(4)=4.999, p=0.008. (D) Left, representative images of c-fos (red) and mCherry projections (green) in the lateral hypothalamus of Nts-cre mice expressing control mCherry (left) or Chr2 (right) virus injected with CNO. n=3, unpaired Students’ t-test, t(4)=5.089, p=0.007. (E) Distance, measured in centimeters (cm), traveled in the elevated plus maze, in control mCherry (black) and Chr2 (red) expressing Nts-cre mice. n=5, unpaired Students’ t-test, t(8)=0.9092, p=0.39. (F) Velocity, measured as the cm per s (cm.s^-1^), in the elevated plus maze, in control mCherry (black) and Chr2 (red) expressing Nts-cre mice. n=5, unpaired Students’ t-test, t(8)=1.291, p=0.233. (G) Time, measured in seconds (s), spent in the center and border of an open field in control mCherry (black) and Chr2 (red) expressing Nts-cre mice. n=5. Two-way ANOVA with post-hoc Bonferroni correction, Subject: F(1,8)=0.051, p=0.827; Border/Center: F(1,8)=18.48, p=0.0026; Interaction: F(1,8)=0.791, p=0.4. Data are represented as mean +- SEM.

**Table 1.** c-Fos expression in the mouse brain after restraint stress.

| **Brain Region** | **c-Fos expression** |
| --- | --- |
| Central Amygdala (CeA) | + |
| Basolateral Amygdala (BLA) | ++ |
| Insular Cortex (IC) | ++ |
| Clastrum (CLA) | +++ |
| Lateral Hypothalamic Area (LHA) | +++ |
| Lateral Habenula (Lhb) | ++ |
| Paraventricular nucleus of the Thalamus (PVT) | ++ |
| Lateral Septum (LS) | +++ |
| Medial Preoptic Nucleus (MPO) | +++ |
| Periqueductal Gray (PAG) | ++ |
| Bed Nucleus of the stria terminalis (BNST) | +++ |

**Score:** - (no expression), + (low expression), ++ (moderate expression), +++ (high expression)

